# The establishment of variant surface glycoprotein monoallelic expression revealed by single-cell RNA-seq of *Trypanosoma brucei* in the tsetse fly salivary glands

**DOI:** 10.1101/2021.03.01.433049

**Authors:** Sebastian Hutchinson, Sophie Foulon, Aline Crouzols, Roberta Menafra, Brice Rotureau, Andrew D. Griffiths, Philippe Bastin

## Abstract

The long and complex *Trypanosoma brucei* development in the tsetse fly vector culminates when parasites gain mammalian infectivity in the salivary glands. A key step in this process is the establishment of monoallelic variant surface glycoprotein (*VSG*) expression and the formation of the VSG coat. The establishment of VSG monoallelic expression is complex and poorly understood, due to the multiple parasite stages present in the salivary glands. Therefore, we sought to further our understanding of this phenomenon by performing single-cell RNA-sequencing (scRNA-seq) on these trypanosome populations. We were able to capture the developmental program of trypanosomes in the salivary glands, identifying populations of epimastigote, gamete, pre-metacyclic and metacyclic cells. Our results show that parasite metabolism is dramatically remodeled during development in the salivary glands, with a shift in transcript abundance from tricarboxylic acid metabolism to glycolytic metabolism. Analysis of *VSG* gene expression in pre-metacyclic and metacyclic cells revealed a dynamic *VSG* gene activation program. Strikingly, we found that pre-metacyclic cells contain transcripts from multiple *VSG* genes, which resolves to singular *VSG* gene expression in mature metacyclic cells. Single molecule RNA fluorescence *in situ* hybridisation (smRNA-FISH) of *VSG* gene expression following *in vitro* metacyclogenesis confirmed this finding. Our data demonstrate that multiple *VSG* genes are transcribed before a single gene is chosen. We propose a transcriptional race model governs the initiation of monoallelic expression.

## Introduction

African trypanosomes are single-cell flagellated parasites that cause Human African Trypanosomiases, and nagana in cattle in sub-Saharan Africa. The parasites have a digenetic life cycle, cycling through a mammalian reservoir host and a tsetse fly vector host. During the developmental phase in the fly, the parasite progresses through multiple stages before acquiring mammalian infectivity in a process called metacyclogenesis in the salivary glands (Rotureau and Van Den Abbeele 2013). Trypanosome cells can be classified into multiple stages grouped in two morphotypes: trypomastigote and epimastigote forms (Figure 1-figure supplement 1), defined by the relative positions of the nuclear and mitochondrial (kinetoplast) DNA within the cell (Hoare and Wallace 1966). Following a tsetse blood meal, the trypomastigote stumpy-form parasites ingested from the blood rapidly differentiate into trypomastigote procyclic form (PCF) parasites which colonise the posterior midgut of the fly. Procyclic trypanosomes elongate and migrate through the alimentary tract to reach the cardia (or proventriculus), while differentiating into epimastigote forms.

**Figure 1.**
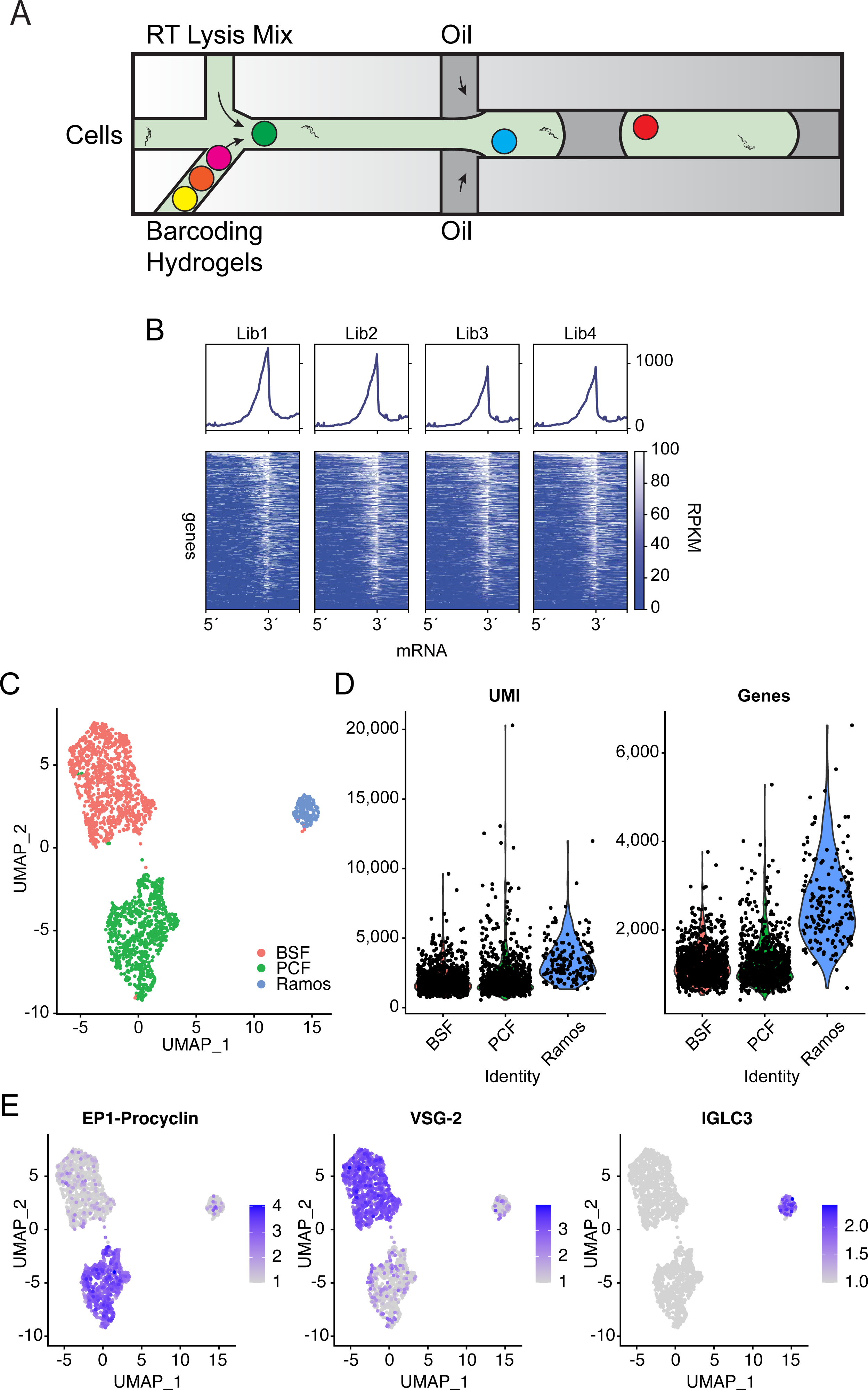
inDrop analysis of cultured bloodstream form and procyclic trypanosomes. A. Schematic of the inDrop microfluidic chip, adapted from Klein, Mazutis et al. (2015). B. inDrop performed in a microfluidic format, using a single primer to replace the BHMs (no single-cell resolution). The plots show the reads per kilobase per million mapped (RPKM) for annotated *T. brucei* transcripts aligned from their 5’splice site to the 3’polyadenylation sequence. Profile plots (above) show the average RPKM across the ∼7,500 non-redundant gene set for trypanosomes. Heatmaps (below) show the RPKM for individual transcripts. C. UMAP projection of single-cell barcoding data for a population of cells containing calculated proportions of 47.5% for each BSF and PCF, and 5% Ramos cells. Measured cell proportions are 51% BSF, 41% PCF and 8% Ramos. D. Violin plots depict the total number of UMIs and genes captured per cell within BSF, PCF and Ramos cell clusters. E. UMAP projections from C overlaid with colour scale of gene expression values for *EP1* procyclin (Tb927.10.10260), *VSG-2* (Tb427.BES40.22) and Immunoglobulin lambda constant 3 (IGLC3).

After a first asymmetric division, epimastigote parasites then migrate to the salivary glands where they attach to the epithelium via their flagellum and start proliferating to colonise it (Van Den Abbeele, Claes et al. 1999, Sharma, Peacock et al. 2008). This population of attached epimastigote cells enters another asymmetric division, producing trypomastigote pre-metacyclic cells which then mature further to produce free mammalian-infective metacyclic cells that fill the lumen of the salivary glands and will be injected to the next host with the tsetse saliva (Rotureau, Subota et al. 2012) (Figure 1-figure supplement 1). The two mitotic cell divisions of attached epimastigote cells, producing either an attached epimastigote or a pre-metacyclic trypomastigote daughter cell occur continuously and in parallel during the entire life of the tsetse, optimizing parasite transmission. An additional heterogeneous parasite population is found in the salivary glands, composed of meiotic cells and gametes (Peacock, Ferris et al. 2011, Peacock, Bailey et al. 2014). These processes lead to a heterogeneous population of cells within the salivary glands. RNA binding proteins appear to be the key regulators underpinning the trypanosome developmental program (Kolev, Ullu et al. 2014). For instance, ALBA3 restricts development in the cardia (Subota, Rotureau et al. 2011) whereas overexpression of RNA binding protein 6 (RBP6) is sufficient to drive differentiation to infective metacyclic cells (Kolev, Ramey-Butler et al. 2012).

In addition to the dramatic remodeling described above, trypanosomes modulate their energy metabolism in response to their specific environments (Smith, Bringaud et al. 2017). In the mammalian host, parasites utilize glucose as a primary carbon source to produce ATP through the glycolytic pathway that is restricted in specialized organelles named glycosomes. In contrast, in the tsetse fly midgut, parasites mostly use proline to feed the mitochondrial tricarboxylic acid (TCA) cycle and oxidative phosphorylation pathways to produce ATP (review in Smith, Bringaud et al. (2017)).

A key step in the metacyclogenesis process is the establishment of monoallelic *VSG* gene expression (Tetley, Turner et al. 1987). African trypanosomes have an exclusively extracellular lifecycle, meaning that in the mammalian host they are continually exposed to the immune system. To evade humoral immune attack, the parasite covers its surface in a monolayer of 10^7^ molecules of a single species of VSG protein which protects from complement mediated lysis (Cross 1975, Mosser and Roberts 1982). It is put under continuous pressure to switch from the adaptive immune system, and the parasite evades this by antigenic variation, a process of VSG switching (Horn 2014).

The epigenetic control of monoallelic expression of *VSG* genes underpins this antigenic variation (Duraisingh and Horn 2016). Transcription of *VSG*s is highly structured; *VSG* genes are always transcribed from a telomere proximal site known as a *VSG-*expression site (*VSG*-ES), by RNA polymerase I (RNA Pol I) (Zomerdijk, Kieft et al. 1991, Gunzl, Bruderer et al. 2003). The genome has 19 bloodstream form *VSG*-ES which are polycistronic transcription units, and 8 monocistronic metacyclic *VSG*-ES (*mVSG*-ES) (Hertz-Fowler, Figueiredo et al. 2008, Muller, Cosentino et al. 2018). In addition, the trypanosome genome encodes several thousand silent *VSG* genes and gene fragments in a sub-telomeric archive which hugely increases the antigen diversity available for antigenic variation (Berriman, Ghedin et al. 2005). The single active *VSG*-ES is transcribed from a sub-nuclear organelle called the expression site body (ESB), an extra-nucleolar RNA Pol I focus (Navarro and Gull 2001). Transcription is initiated at all *VSG-*ES, however only elongates over a single *VSG-*ES (Kassem, Pays et al. 2014). Maintenance of monoallelic *VSG* expression depends on the *VSG* Exclusion complex (VEX) complex, which associates with the ESB (Glover, Hutchinson et al. 2016, Faria, Glover et al. 2019), and an mRNA trans-splicing locus, forming a transcription and splicing factory (Faria, Luzak et al. 2021). In addition, several chromatin factors, including the sheltrin component RAP1 and the histone chaperone CAF1 have been implicated in maintenance of monoallelic expression (Yang, Figueiredo et al. 2009, Alsford and Horn 2012, Faria, Glover et al. 2019).

The establishment of monoallelic expression in metacyclic trypanosomes is relatively poorly understood. Immunogold staining of salivary gland trypanosomes and double immunofluorescence staining of *in vitro* derived metacyclic parasites demonstrates that monoallelic expression is established in the salivary glands of the tsetse fly (Tetley, Turner et al. 1987, Ramey-Butler, Ullu et al. 2015). In bloodstream parasites, monoallelic expression can undergo transcriptional switching events, where the active *VSG*-ES is switched off and a previously silent one is activated epigenetically (Horn 2014), requiring the re-establishment of monoallelic expression. Analysis of BSF cells following a forced *VSG*-ES switch showed that transcription transiently increased at silent *VSG-*ES prior to switching the active *VSG*-ES. This was proposed as “probing” of silent *VSG*-ES before commitment to switching (Aresta-Branco, Pimenta et al. 2016). Only one factor has so-far been linked to the establishment of monoallelic expression; the histone methyltransferase DOT1b trimethylates Histone H3 lysine 76 (Janzen, Hake et al. 2006) and is required to switch off a previously active *VSG-*ES following the establishment of a new active site (Figueiredo, Janzen et al. 2008).

The limiting nature and diverse cellular composition of salivary gland trypanosomes has made these parasites less experimentally tractable than *in vitro* systems. These parasites are not amenable to axenic culture (metacyclic cells are G0 arrested), and genetic manipulation tools are not as well established as in cultured parasites. We therefore applied a single-cell RNA-sequencing (scRNA-seq) approach to salivary gland derived trypanosomes, which overcomes many experimental boundaries through increased sensitivity and granularity to address the lack of resolution in these cell types. Here, we dissect the architecture of parasite development as they acquire mammalian infectivity in the salivary glands using inDrop. InDrop is a scRNA-seq technology wherein single cells are co-encapsulated inside nanoliter droplets with single barcoded hydrogel microspheres (BHMs) coupled to barcoded cDNA primers that hybridize to the 3’ poly(A) tail of mRNA (Figure 1A), allowing sequencing of several thousand cells per experiment. Reverse transcription of mRNAs from each cell generates cDNAs tagged with a unique barcode, specific to each cell, permitting the informatic decomposition of single-cell transcriptomes (Klein, Mazutis et al. 2015, Zilionis, Nainys et al. 2017).

In this study, we first demonstrated that trypanosomes are compatible with 3’ mRNA barcoding using inDrop BHMs. After having generated single cell transcriptomes for cultured bloodstream (BSF) and PCF trypanosomes, we performed inDrop with parasites from the salivary glands. This revealed the presence of 4 cell clusters in the salivary glands, corresponding to 1) attached epimastigote cells, 2) gametes and meiotic cells, 3) pre-metacyclic trypomastigote cells and 4) metacyclic parasites. We delineated transcriptomic changes to energy metabolism pathways, revealing a coherent developmental program in this organ towards infectivity. Finally, we have performed the first analysis of the establishment of monoallelic VSG expression *in vivo*. We show that the establishment of monoallelic expression is a two-step process, where pre-metacyclic cells first initiate expression of multiple *mVSG* transcripts, followed by monoallelic expression in metacyclic parasites. We further validated this finding by an orthogonal method, employing smRNA-FISH with parasites undergoing metacyclogenesis *in vitro*.

## Results

### Assessment of single-cell RNA-seq using inDrop applied to *Trypanosoma brucei*

To test whether trypanosomes would not be visibly stressed by the inDrop pipeline, we monitored the survival and RNA metabolism using ALBA3 and DHH1 as markers for cellular stress in procyclic form (PCF) cells, in conditions which mimic the inDrop protocol. Following starvation in PBS for 2 h at 27°C, ALBA3 and DHH1 form RNA granules in the cytoplasm (Subota, Rotureau et al. 2011), however when cells are placed on ice for 2h in phosphate buffered saline, no RNA granule formation was observed (Figure 1-figure supplement 2A). In addition, cells remained alive based on their motility after returning them to 27°C SDM-79 medium (97 % untreated and 94 % 2h PBS) (Figure 1-figure supplement 2B), indicating that they are not visibly stressed in the conditions required for inDrop.

We first characterized the performance of the inDrop pipeline with cultured Antat1.1E PCF trypanosomes expressing EP (Glu-Pro) / GPEET (Gly-Pro-Glu-Glu-Thr) repeat containing procyclins. *T. brucei* mRNAs are poly-adenylated and therefore compatible with priming of cDNA synthesis using poly(T)VN, so we began by performing ‘bulk’ experiments using a primer containing a T7 promoter for cDNA amplification by *in vitro* transcription (IVT), an Illumina sequencing adaptor, a single inDrop cell barcode, unique molecular identifiers UMI (Islam, Zeisel et al. 2014) and a poly(T)VN sequence for mRNA capture (see Materials and Methods). Library sequencing revealed that the majority of barcoded cDNAs corresponded to the annotated 3’-end of transcripts, as expected (Figure 1B). We therefore proceeded to single cell barcoding of trypanosomes. We mixed cultured BSF and PCF trypanosomes (47.5 % each) with Ramos cells (5%), a human B-cell line derived from Burkitt’s lymphoma (Klein, Giovanella et al. 1975) as a positive control. The inclusion of Ramos cells also allows us to estimate the amount of ambient RNA within the cell suspension which is detected by inDrop. We constructed a sequencing library, and generated 77.4 million reads. To facilitate the use of trypanosome genomic sequences for which no reference strain is available, we developed a bespoke analysis pipeline to create a count matrix (see Materials and Methods). After processing the reads with this pipeline, we used the Seurat and single cell transform (SCT) R packages for data analysis (Butler, Hoffman et al. 2018, Hafemeister and Satija 2019, Stuart, Butler et al. 2019) and determined that the transcriptomes of 1,979 cells were captured, with an average of 2,195 UMIs and 1,347 genes per cell. The emulsion contained approximately 4,900 cells, suggesting that we recovered approximately 40 % of the cells through inDrop. Dimensionality reduction using the Uniform Manifold Approximation and Projection (UMAP) algorithm (L McInnes, J Healy et al. 2018) revealed three clusters which corresponded to the three expected cell types in approximately the expected proportions (Figure 1C), with 164 (8%) Ramos cells, 809 (41%) procyclic cells and 1,006 (51%) BSF cells. Within clusters, we detected an average of 1,932 UMIs and 1,211 genes for BSF, 2,268 UMIs and 1,258 genes for PCF cells and an average of 3,447 UMIs and 2,627 genes in Ramos cells (Figure 1D). Each trypanosome cell has between 20,000-40,000 mRNA molecules per cell (Haanstra, Stewart et al. 2008) therefore these data are consistent with an mRNA capture efficiency between 4.8 % and 11.3 %, similar to the 7.1 % reported by Klein, Mazutis et al. (2015). We next plotted percentage of human and trypanosome UMIs for each barcode (Figure 1-figure supplement 2C). This revealed that human barcodes contained higher proportions of trypanosome UMIs than the converse. We hypothesized that this was potentially due to trypanosome cell fragility which leads to release of trypanosome mRNAs from lysed cells. We therefore modified our protocol to maintain cell suspensions on ice until the moment they are encapsulated (see Materials and Methods).

Marker gene analysis for each cluster revealed significantly differentially expressed genes, corresponding to the known biology of each respective cell type. In the BSF cluster, marker genes included surface proteins such as the active *VSG-2* (Tb427.BES40.22) (Figure 1E), *GRESAG 4* (Tb927.7.6080), *ISG65* (Tb927.2.3270), and *GPI-PLC* (Tb927.2.6000), as well as transcripts involved in the glycolytic pathway, such as *AOX* (Tb927.10.7090), *ALD* (Tb927.10.5620), and *PYK1* (Tb927.10.14140), for example. The PCF cluster markers included the procyclic surface markers *EP1* (Tb927.10.10260) (Figure 1E), *EP2* (Tb927.10.10250), *GPEET* (Tb927.6.510) and *PARP A/EP3-2* (Tb927.6.450), the electron transport chain component *COXIV* (Tb927.1.4100), an amino acid transporter (*AAT10-2*) (Tb927.8.8300), the RNA binding protein *UBP2* (Tb927.11.510) and *HSP90* (Tb927.10.10980), all of which are seen to be up-regulated in PCF using bulk RNA-seq (Siegel, Hekstra et al. 2010). In the Ramos cluster, markers included genes such as immunoglobulin lambda constant 3 (*IGLC3*) (Figure 1E) and prothymyosin alpha (*PTMA*), as expected. These data correspond with known biology from bulk transcriptomics experiments and therefore demonstrate that inDrop is capable of single-cell analysis of trypanosome transcripts.

### Single-cell analysis of trypanosome salivary gland transcriptomes using inDrop

We next proceeded to single-cell barcoding of trypanosomes from tsetse fly salivary glands. We infected cages of 50 teneral (1-2 days post-eclosion) male tsetse flies with *in vitro* generated stumpy-form Antat1.1E (EATRO1125) *T. b. brucei* and recovered salivary gland parasites by dissection 28 days post infection. We obtained 4.6 x10^4^ parasites from a total of 72 dissected flies (Figure 2-figure supplement 1). We then generated and sequenced single-cell barcoded libraries (Figure 2-figure supplement 2A), and developed an optimized bioinformatic pipeline for *T. brucei*, recovering data for 2,279 cells from two technical replicates (Figure 2 and Figure 2-figure supplement 2B), detecting an average of 959 UMIs per replicate (Figure 2-supplement 2B). As no genome sequence is available for the Antat1.1 strain used here, we identified *VSG* genes expressed in the salivary glands by the Antat1.1 strain by aligning reads to the *VSG*nome for Antat1.1E (EATRO1125) (Cross 2017). We then selected the top 8 sequences, in line with the reported number of *mVSG*-ES in the Lister-427 strain (Muller, Cosentino et al. 2018).

**Figure 2:**
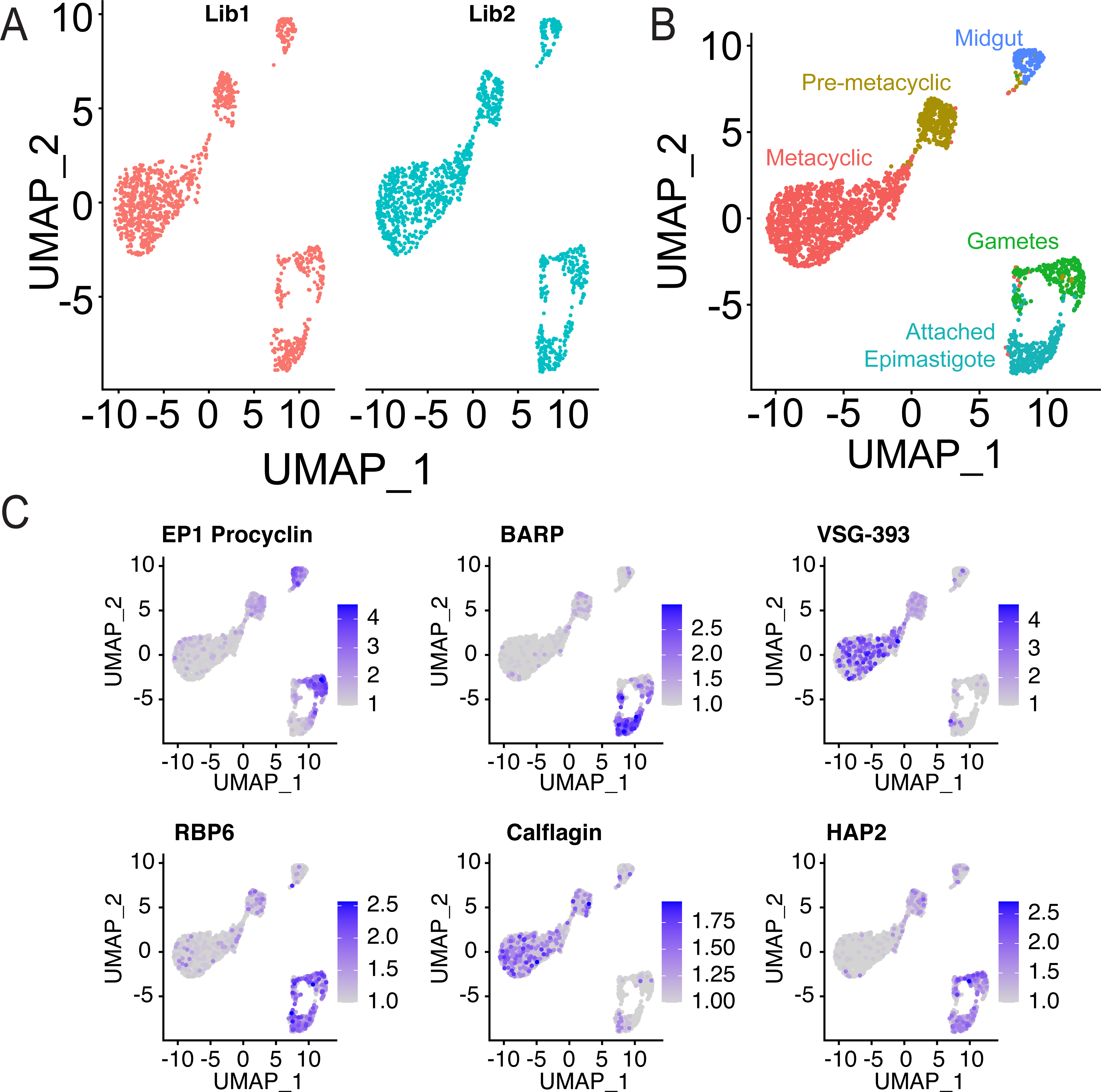
InDrop barcoding of salivary gland *T. brucei*. A. UMAP projection of two technical replicates (Lib1 and Lib2, red and blue, respectively). B. Merged UMAP projection of technical replicates with clustering information as indicated. C. Gene expression levels for marker genes in the salivary glands, *EP1 procyclin* (midgut stages) (Roditi 1987 nature): Tb927.10.10260, *BARP* (epimastigote stage): Tb927.9.15640 (Urwyler, Studer et al. 2007), *VSG-393*: (Genbank) KC612418.1 (Cross 2017), *RBP6*: Tb927.3.2930 (Kolev, Ramey-Butler et al. 2012), *Calflagin*: Tb927.8.5440 (Rotureau, Subota et al. 2012), *HAP2*: Tb927.10.10770 (Fedry, Liu et al. 2017).

We used the Seurat R package to analyze our technical replicates (Butler, Hoffman et al. 2018, Hafemeister and Satija 2019). Count matrices were compiled from all aligned reads, except for *VSG* genes, where we used a stringent mapping quality (MapQ40), to avoid mis-aligned reads being spuriously counted. We next examined several technical aspects of inDrop, Firstly, we examined the background from ambient RNA by looking at the detection rate in a cluster of Ramos cells, which were incorporated into our experiment. These data showed that we did not detect more than 2 *mVSG* UMIs per barcode in Ramos cells (Figure 2-figure supplement 3A). We therefore applied a >2 UMI cutoff for our *<u>m</u>VSG* analyses below. We established the percentage of stringently aligned reads (mapping quality >40) in our data. This showed improvement over our *in vitro* data, with two distinct clusters of cells for human and trypanosome UMIs (Figure 2-figure supplement 3B). We next considered whether sequencing errors could contribute to barcode switching. Our bioinformatics pipeline permits up-to two errors per cell barcode, so we compared the edit distance between all cell barcodes in our study. The mean editing distance between barcodes was 11.4 nt, and only 0.017 % of barcode pairs were within an editing distance of 2 (Figure 2-figure supplement 3C). These parameters allowed us to effectively estimate the noise for our downstream analyses.

Following data preprocessing and integration of the technical replicates, we visualized the datasets using UMAP projections (L McInnes, J Healy et al. 2018) (Figure 2A). Clustering identified 5 clusters (Figure 2B), whose developmental state could be identified using known marker genes (Figure 1E). *EP1 Procyclin* (Tb927.10.10260) is a surface protein found in midgut forms (Roditi, Carrington et al. 1987), and its transcript can also be detected in salivary gland parasites (Urwyler, Studer et al. 2007). Brucei alanine-rich protein (*BARP*, Tb927.9.15640) is a surface protein present in attached epimastigote parasites (Urwyler, Studer et al. 2007). Metacyclic VSG expression is a marker for metacyclic cells (Rotureau, Subota et al. 2012), and *VSG-393* is the most abundant *VSG* transcript expressed in our dataset. RBP6 (Tb927.3.2930) is an RNA binding protein which drives developmental progression to infectivity in salivary glands, and is found in epimastigote forms in the salivary glands (Kolev, Ramey-Butler et al. 2012, Vigneron, O’Neill et al. 2020). Calflagin is a flagellar calcium binding protein whose expression is up-regulated in pre-metacyclic and metacyclic forms (Rotureau, Subota et al. 2012). Finally, HAP2 (Tb927.10.10770) is a membrane-membrane fusion protein found conserved in nature and expressed in the gametes of many species (Figure 2C) (Fedry, Liu et al. 2017). Further evidence from bulk transcriptomics of midgut, cardia (proventriculus) or salivary gland tissues (Kolev, Ramey-Butler et al. 2012) were used to identify midgut forms. Of the 226 marker genes identified for the midgut forms cluster in our inDrop data, 82.3 % of genes were maximally expressed in the midgut samples of bulk transcriptomics (Kolev, Ramey-Butler et al. 2012), further supporting this annotation. In total, we identified 142 midgut forms, 296 attached epimastigote cells, 280 gametes, 317 pre-metacyclic and 1,244 metacyclic cells. These data indicate that we are able to capture single cell transcriptomes of these salivary gland parasites, and identify cell states from these transcriptomes, using inDrop.

### Metabolic transcript analysis reveals dramatic remodeling during metacyclogenesis

To further our understanding of developmental changes in the salivary glands, we assessed the expression of glycolysis and TCA cycle genes expressed in the transcriptomes of parasites from the salivary glands. Our analysis reveals that parasites remodel the expression of metabolic enzymes during differentiation to metacyclic cells (Figure 3, and Figure 3-figure supplement 1). We were able to assess the expression changes for 12 glycolysis enzyme transcripts and 7 TCA cycle transcripts (Figure 3A). We observed a common expression profile for most genes at the cluster (Figure 3B and Figure 3-figure supplement 1) and single cell level (Figure 3C). Glycolytic enzymes increased in their expression between pre-metacyclic and metacyclic clusters, with hexokinase (*HK1*, Tb927.10.2010) and ATP-dependent 6-phosphofructokinase (*PFK*, Tb927.3.3270) being clear examples of this. We observed a decline in TCA cycle enzyme expression during metacyclogenesis, with pre-metacyclic cells positioned as a clear intermediate stage between epimastigote forms (Attached and Gametes) and metacyclic cells. For instance, aconitase (*ACO*, Tb927.10.14000) and glycosomal isocitrate dehydrogenase (*IDH*, Tb927.11.900) both decrease from their peak expression in attached epimastigote cells to their lowest in metacyclic parasites with pre-metacyclic cells as intermediates. This indicates that the data clustering is indeed reflecting the true developmental progression of these cells, consistent with previous electron microscopy observations on glycosomal and mitochondrial remodeling during this development (Tetley and Vickerman 1985, Kolev, Ramey-Butler et al. 2012).

**Figure 3:**
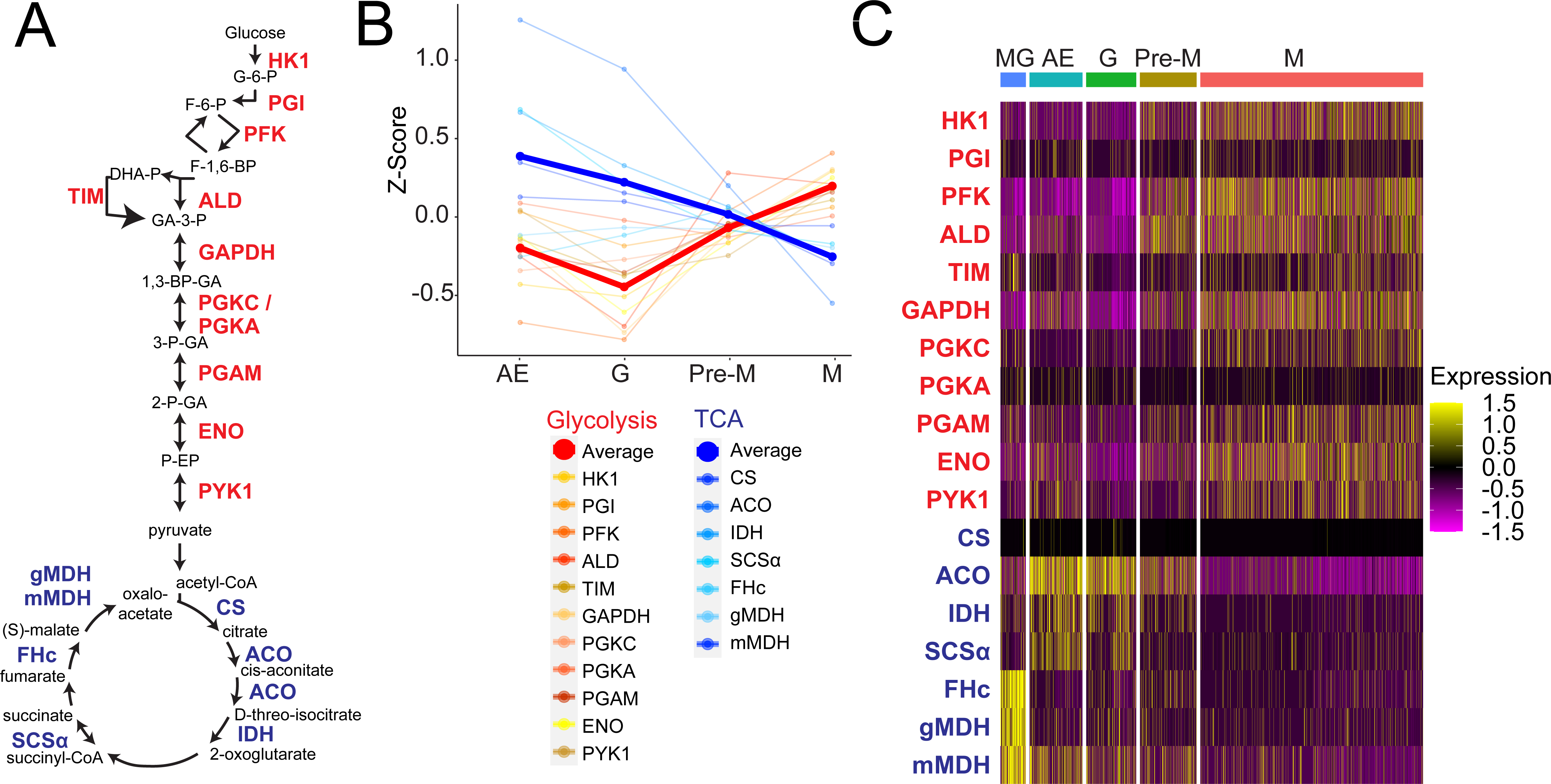
Cluster-based analysis of main energy metabolism transcripts. A. The schematic shows the glycolytic and TCA pathways (Shameer, Logan-Klumpler et al. 2015). Compounds (black text) and enzymes (red text for glycolysis, blue text for TCA cycle) are shown and reactions are represented by arrows. B. Plot shows the expression Z-scores for each glycolysis or TCA cycle enzyme transcript retained after SCT normalization (Hafemeister and Satija 2019). Thicker lines show the average across each cohort. AE: attached epimastigote cells; G: Gamete cells; Pre-M: pre-metacyclic cells: M: metacyclic cells. C. Single cell level analysis of metabolic transcripts. Cells are arranged by cluster (left to right) and relative transcript expression shown per cell. Colours represent Z-transformed expression values. Cluster abbreviations as for B. Abbreviations of metabolites: G-6-P: glucose-6-phosphate; F-6-P: fructose-6-phosphate; F-1,6-BP: fructose-1,6-bisphosphate; DHA-P: dihydroxyacetone phosphate; GA-3-P: glyceraldehyde-3-phosphate; 1,3-BP-GA: 1,3-bisphosphoglycerate; 3-P-GA: 3-phosphoglycerate; 2-P-GA: 2-phosphoglycerate; P-EP: phospho-enol pyruvate. Abbreviations of enzymes: HK1: hexokinase 1; PGI phosphoglucose isomerase; PFK: phosphofructokinase; ALD: aldolase; TIM: triose-phosphate isomerase; GAPDH: glyceraldehyde-3-phosphate dehydrogenase; PGK(C/A): phosphoglycerate kinase, PGAM: phosphoglycerate mutase; ENO: enolase; PYK1: pyruvate kinase; CS: citrate synthase; ACO: aconitase; IDH: isocitrate dehydrogenase; SCS*α*: succinyl coenzyme A synthetase; FHc, fumarate hydratase, cytosolic; gMDH: glycosomal malate dehydrogenase; mMDH: mitochondrial malate dehydrogenase.

### Identification of a putative gamete cluster

A subpopulation of trypanosomes undergoes meiosis I and II in the salivary glands, producing gamete cells that can recombine sexually (Peacock, Ferris et al. 2011). *HAP2* expression has not been previously profiled during parasite development in the salivary glands. In our data, expression of *HAP2* was significantly upregulated in gamete cells compared to the other clusters (average log2 FC 0.74, detected in 82.5 % gamete cells and 36.6 % cells in other clusters, adjusted p.val = 0. 9.75 x 10^-50^, negative binomial test, Supplementary table 1). Additionally, and unexpectedly, *BARP* expression was significantly reduced in these cells compared to gamete cells (average log2 FC −0.83, detected in 94.6 % attached epimastigote cells and 61.1 % gamete cells, adjusted p.val = 9.68 x 10^-36^, negative binomial test) and *EP1 procyclin* expression up-regulated (average log2 FC 1.45, detected in 58.1% and 87.5% of attached epimastigote cells and gamete cells, respectively, adjusted p.val = 3.26 x 10^-56^), potentially explaining the presence of these transcripts in previous studies (Urwyler, Studer et al. 2007). We also detected a significant up-regulation of the gametocytogenesis marker *HOP1* (Tb927.10.5490) (Peacock, Ferris et al. 2011) in this cluster compared to the other clusters (average log2 FC 0.80, detected in 7.5 % gamete cells and 1.8 % cells in other clusters, adjusted p.val = 1.54 x 10^-4^). These data therefore support the assignment of gamete cells to this cluster.

Attached epimastigote cells undergo asymmetrical division to produce one attached epimastigote daughter and one pre-metacyclic trypomastigote daughter (Rotureau, Subota et al. 2012). We examined these cells in more detail (Figure 4A). We searched for cells expressing the histones (*H1* Tb927.11.1800, *H2A* Tb927.7.2820, *H2B* Tb927.10.10460, *H3* Tb927.1.2430 and *H4* Tb927.5.4170), as a proxy for S-phase cells (Ersfeld, Docherty et al. 1996), and discovered two sub-clusters of cells expressing these transcripts (Figure 4B). To investigate this more quantitatively, we sub-divided the S-Phase cluster and assessed the marker genes expressed in these clusters (Figure 4C). This revealed that *BARP* and the set of 5 canonical histones were significantly associated with the “BARP-S” cluster. *HAP2* was significantly associated with both clusters, albeit more strongly with the HAP2-S cluster (Supplementary table 2). The presence of these two clusters raises the questions as to whether there are two potential modes of development in the salivary glands.

**Figure 4:**
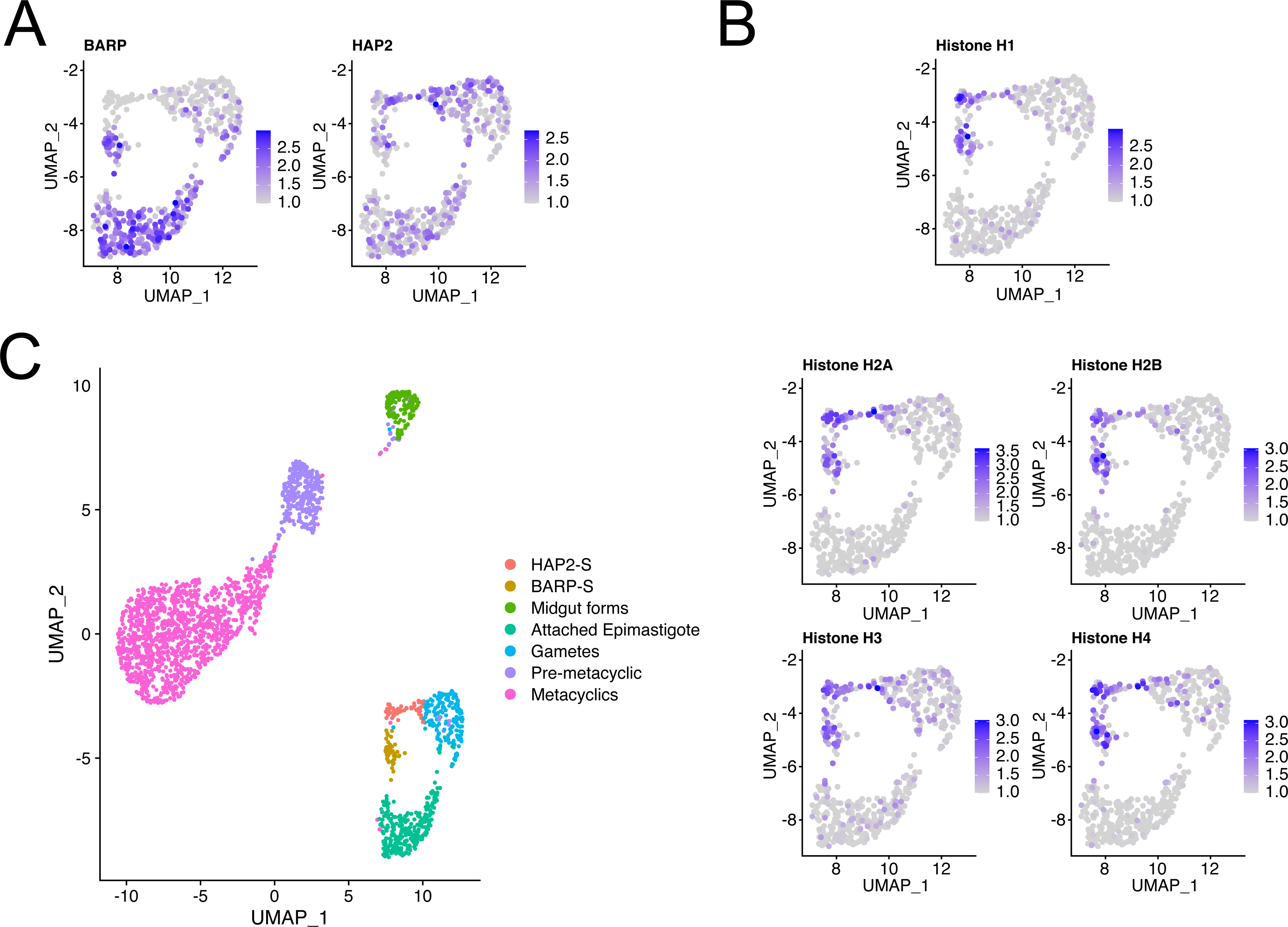
Analysis of gamete cluster. A. Normalised expression values for *BARP* (Tb927.9.15640) and *HAP2* (Tb927.10.10770) overlaid on UMAP projections of attached epimastigote and gamete cell clusters. B. Normalised expression values for canonical histones: H1 Tb927.11.1800, H2A Tb927.7.2820, H2B Tb927.10.10460, H3 Tb927.1.2430 and H4 Tb927.5.4170. C. Sub-clustering of cells in S-Phase. Clusters were manually re-annotated using the Seurat function CellSelector. Two new clusters of cells expressing both core histones and either BARP or HAP2 were created and are indicated in brown and red, respectively.

### Establishment of *mVSG* monoallelic expression

VSG protein expression is initiated during pre-metacyclic to metacyclic differentiation (Tetley and Vickerman 1985, Rotureau, Subota et al. 2012). Consistent with this, our data revealed, as expected, the presence of very few *mVSG* UMIs in attached epimastigote cells with 85% of cells containing no detectable *mVSG* transcripts, compared to pre-metacyclic and metacyclic clusters (Figure 5-figure supplement 1). Using our Ramos cell data to estimate the amount of ambient RNA in the cell suspension detected by inDrop, we found that attached epimastigote cells contained more *mVSG* UMIs on average than Ramos cells, where these genes are not present (3.36 vs 0.19 UMIs per cell, respectively), suggesting that there is some transcription of these loci, consistent with ubiquitous transcription initiation from RNA-Pol I promoters (Kassem, Pays et al. 2014). We observed an increase in the average number of *mVSG* UMIs per cell as parasites proceeded through their developmental program (Figure 5-figure supplement 1), increasing from a mean of 3.36 UMIs in attached epimastigote cells to 10.0 UMIs and 21.7 UMIs in pre-metacyclic and metacyclic cells, respectively.

**Figure 5:**
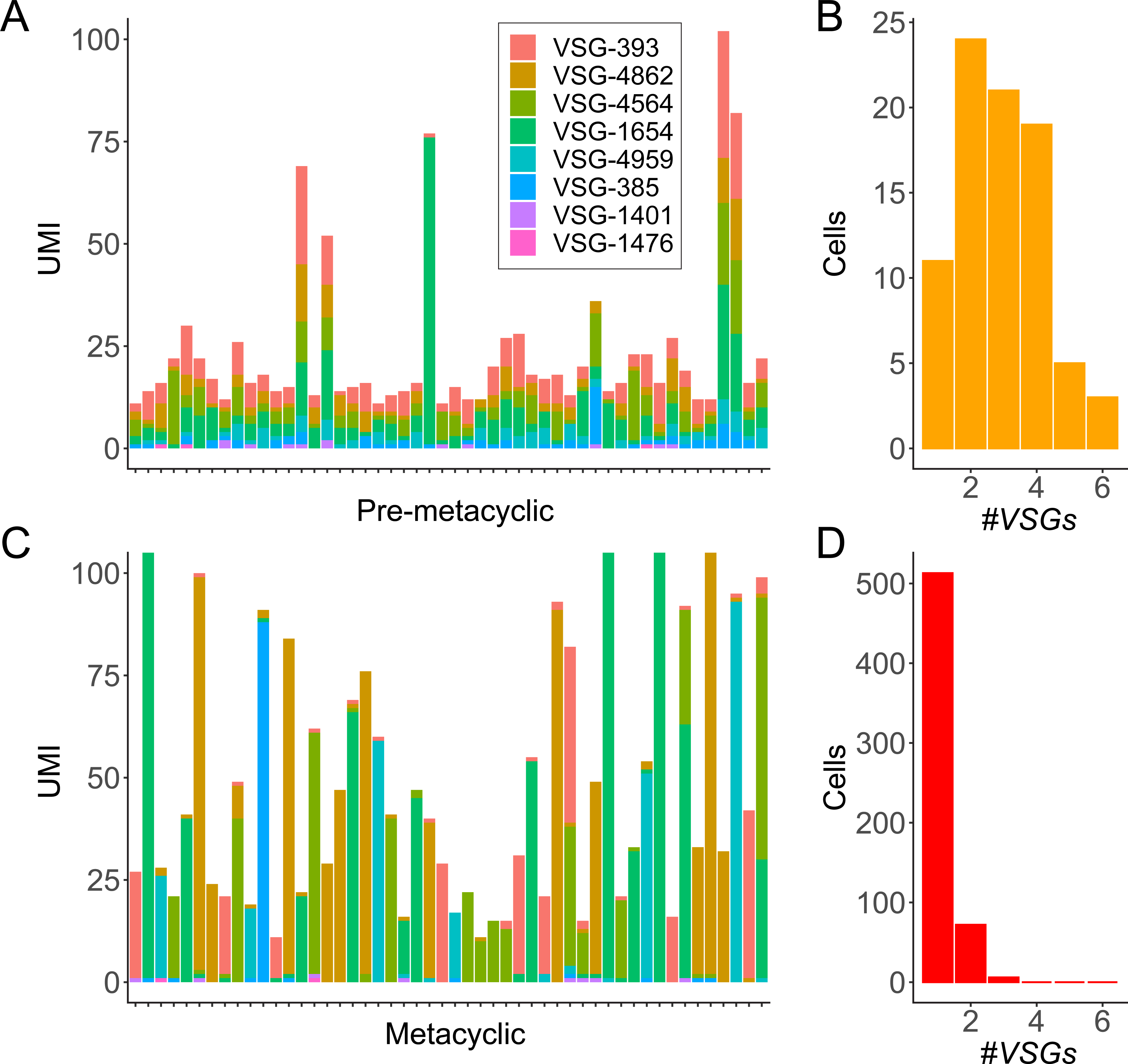
Developmental progression of *VSG* expression. A. *VSG* expression data for a random subset of 50 pre-metacyclic cells, with a total *VSG* UMI count > 10. Data are raw (unscaled) UMI counts. Each column represents a cell and each colour a different *VSG*. B. Histogram showing the number of *VSG* expressed for all pre-metacyclic cells (per *VSG* UMI count > 2, all cells in cluster). C. As for A except that the data are a subset of 50 metacyclic cells with total *VSG* UMI counts >10. D. Histogram as for B except showing metacyclic *VSG* transcript diversity (per *VSG* UMI count >2, all cells in cluster).

We next assessed the diversity of *mVSG* expression during the differentiation process. Plotting the expression profile of individual pre-metacyclic cells showed that the majority of cells in this cluster express multiple different *mVSG* genes (Figure 5A). We quantified the number of different *mVSG* genes expressed per cell (i.e. the diversity of UMIs per cell) in cells with more than 10 *mVSG* UMIs. This revealed, strikingly, that pre-metacyclic cells expressed up-to six different *mVSG* genes. Multi-*mVSG* gene expression (UMI cutoff >2) was observed in 86.7 % of cells, whereas only 13.3 % expressed a single *mVSG* gene. (Figure 5B). Following differentiation to metacyclic form cells, however, *mVSG* expression had resolved into a monoallelic state (Figure 5C), where a single *mVSG* gene was detected in 86.8 % of cells, 12.2 % contained two *mVSG* genes and only 1 % expressed 3 genes types (Figure 5D).

We queried whether the detection of multiple *mVSG* genes in pre-metacyclic cells was a function of sequencing depth for those barcodes, compared to metacyclic cells. We plotted the number of *VSG* detected per barcode against the number of aligned reads, stratified by cluster (Figure 5-figure supplement 2). This revealed that rather than having higher aligned read counts, the median aligned reads per barcode associated to pre-metacyclic cells was lower than metacyclic cells, even when comparing metacyclic cells with a single expressed *mVSG* to pre-metacyclic cells expressing 6 *mVSG* genes (2.2 x 10^4^ vs 1.6 x 10^4^ median aligned reads, respectively). These data show that it is highly unlikely that the detection of multiple *mVSG* transcripts per cell is artefactual due to sequencing depth.

We sought to further validate our findings by turning to previously published scRNA-seq data of salivary gland trypanosomes using both a different trypanosome strain; RUMP503, and a different technique, the 10xGenomics single-cell RNA-seq system (Vigneron, O’Neill et al. 2020). As the complete set of *mVSG* genes was not available for this strain, we used Trinity (Grabherr, Haas et al. 2011) to assemble *mVSG* transcripts *de novo* from bulk RNA-seq data (Savage, Kolev et al. 2016), and retained the 8 most abundant *mVSG* genes for downstream analysis. This allowed us to perform a detailed analysis of *mVSG* expression in these data, which was not possible previously. We reprocessed the scRNA-seq data set using our pipeline, observing similar clusters to Vigneron, O’Neill et al. (2020) (Figure 5-figure supplement 3A,B). We identified a cluster of pre-metacyclic cells, in which we detected multi-*mVSG* gene expression. In cells with more than 10 *mVSG* UMIs, 71 % contained transcripts from multiple *VSG* genes (12 cells), compared to 29 % with one *mVSG* detected (UMI cutoff >2) (Figure 5-figure supplement 3C). In the metacyclic cell clusters, a single *mVSG* gene was detected in 78 % of cells, and 22 % of cells expressed two or more *mVSG* genes, with the majority (86 %) of this subset expressing 2 *mVSG* transcripts. This new finding is similar to our inDrop data, where 87 % of pre-metacyclic cells expressed more than a single *mVSG*. These new analyses uncover that expression of multiple *mVSGs* is a common precursor to monoallelic expression in different trypanosome strains *in vivo*.

To formally demonstrate the existence of cells expressing more than a single *VSG* gene, we performed smRNA-FISH following *in vitro* metacyclogenesis by overexpressing RBP6. This *in vitro* process models the establishment of monoallelic mVSG expression in a third strain, the Lister 427 genetic background. Induction of RBP6 in PCF forms triggers differentiation first to epimastigote forms, which then further differentiate to trypomastigote forms, producing infective metacyclic forms which express a single mVSG (Kolev, Ramey-Butler et al. 2012, Ramey-Butler, Ullu et al. 2015). This process therefore allows us to model the initiation of *mVSG* expression.

We overexpressed RBP6 in PCF cells using a tetracycline inducible system (Trenaman, Glover et al. 2019), and sampled cultures after 3, 4 or 5 days of induction. After probing for *VSG-397*, *VSG-531* and alpha tubulin as a positive control, we detected *VSG* transcripts in ∼5 % of cells (303 cells positive for *VSG* out of a total of 6,452). Within this population, we established 3 categories of cells: “VSG+”, where transcript numbers were quantifiable (193 cells), “VSG++” where transcript numbers were too high to be quantified accurately (but where individual transcripts were discernable) (18 cells), and “VSG+++”, where individual transcripts were no longer discernable (92 cells) (for representative images see Figure 6A, Supplementary table 3).

**Figure 6.**
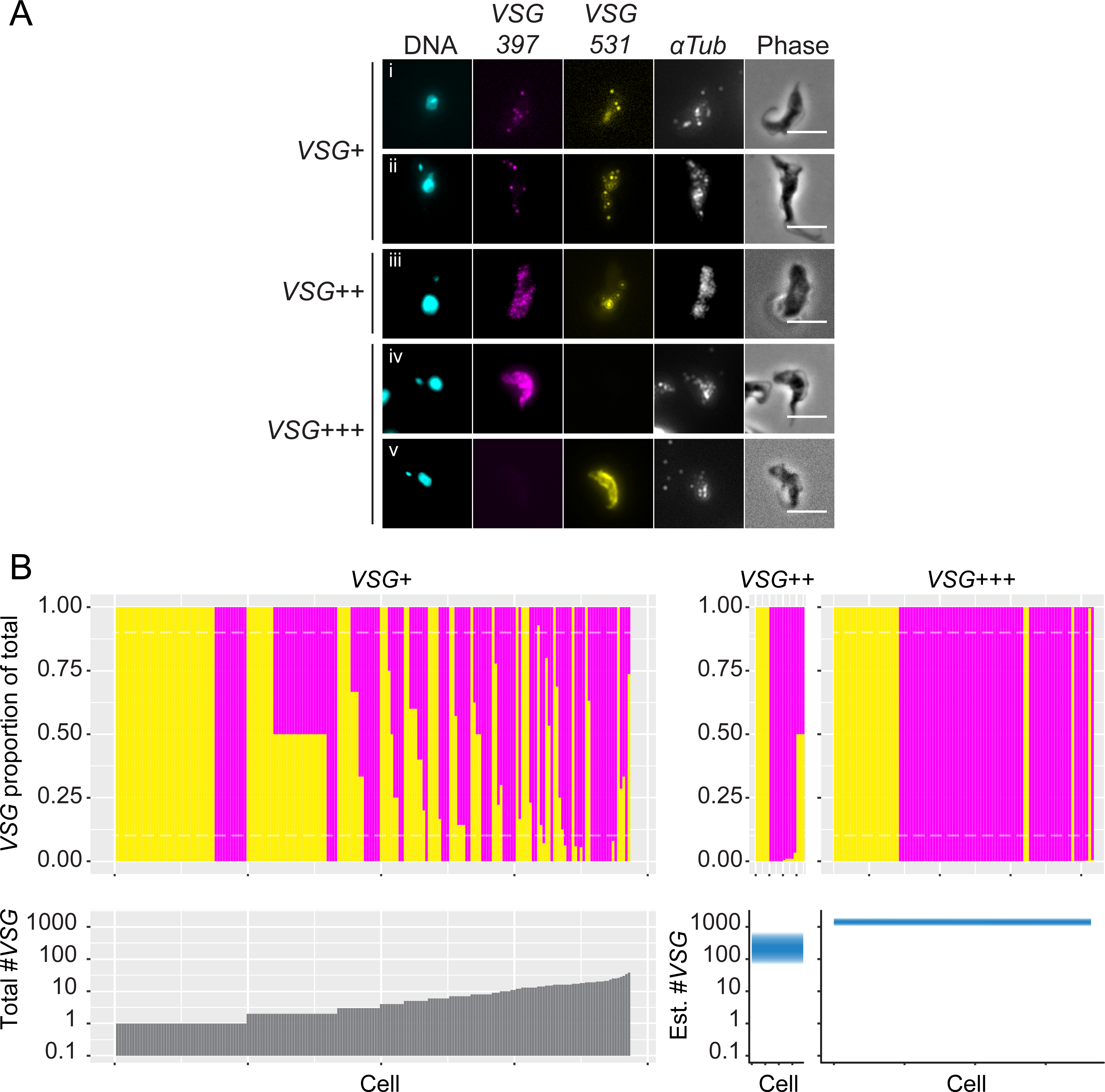
Single molecule RNA-FISH of *in vitro* metacyclogenesis. A. Panel of representative images of VSG+ and VSG++ cells containing both *VSG-397* and *VSG-531* transcripts (*i-iii*) and VSG+++ cells where a single VSG type is present (*iv* and *v)*. B. Above, proportions of *VSG-397* and *VSG-531* transcripts for VSG+, VSG++ and VSG+++ cells. Each cell is plotted as a single column on the x-axis. An estimate of transcript abundance for VSG++ and VSG+++ cells was used based on published estimates of *VSG* transcript abundance in BSF (Haanstra, Stewart et al. 2008). Below, transcript abundance for VSG+ cells, and estimated transcript abundance for VSG+ and VSG++ cells.

To visualize the proportions of *VSG-397* and *VSG-531* transcripts per cell, we estimated the total number of *mVSG* transripts per cell in VSG++ and VSG+++ categories. BSF cells contain on average 2,000 *VSG* transcripts (Haanstra, Stewart et al. 2008). Hence, we set the number of transcripts in VSG+++ cells at 2,000, and VSG++ cells at 200 (an order of magnitude between VSG+ and VSG++) per channel (i.e. if both *VSG-397* and *VSG-531* were “VSG++”, we estimate that cell contains 400 *VSG* transcripts). Cells were therefore classified as VSG+ with less than 200 transcripts, VSG++ with between 200 and 400 transcripts, and VSG+++ with 2000 transcripts or more. We then assessed singular VSG gene expression in each cell category: 30 % (58 cells) VSG+ cells contained transcripts from both *VSG-397* and *VSG-531,* directly demonstrating the presence of cells expressing two different *VSG* genes. This proportion dropped to only 17 % (3 cells) among VSG++ cells and 0 % in VSG+++ cells, indicating that monoallelic expression is established as cells express a larger amount of *mVSG* transcripts (Figure 6B).

## Discussion

Our data demonstrate that the inDrop scRNA-seq approach is compatible with investigation of gene expression in trypanosomes. Despite significant biological constraints (limited access to multiple populations of parasites in low numbers), we were able to delineate the developmental progression of these parasites in the salivary glands, including the identification of a gamete cluster. In our hands, we detected similar numbers of trypanosome UMIs (∼2,000 for cultured cells) compared to Ramos cells (∼3,500), despite having a much lower mRNA content per cell (10^5^ vs 10^4^ for human and trypanosome cells, respectively) (Haanstra, Stewart et al. 2008).

The gamete cell cluster was characterized by a significant upregulation of *HAP2*, a conserved membrane fusion protein required for gamete fusion in diverse organisms (Fedry, Liu et al. 2017), and *HOP1*, a component of synaptonemal complexes formed in the first meiotic division (Peacock, Ferris et al. 2011). We did not, however, observe statistically significant changes in other known marker genes of gametocytogenesis such as *SPO11*, *DMC1* or *MND1* (Peacock, Ferris et al. 2011), likely due the sensitivity of the inDrop. Our analysis of the metabolic transcriptome remodeling indicated that energy metabolism in attached epimastigote cells is dependent on the production of ATP in the mitochondrion, consistent with recent evidence from *in vitro* derived epimastigote parasites using the RBP6 overexpression system (Dolezelova, Kunzova et al. 2020). The most dramatic transcriptome remodeling was following asymmetrical division of attached epimastigote, with pre-metacyclic cells as transition intermediates in terms of metabolic remodeling (Figure 3). These data are consistent with electron microscopy evidence showing an increase in the number and size of glycosomes together with the reduction of mitochondrial branching in metacyclic cells compared to attached epimastigote parasites (Tetley and Vickerman 1985, Kolev, Ramey-Butler et al. 2012).

Our analysis of the establishment of monoallelic *mVSG* expression has uncovered that this is a phased process, first involving the activation of multiple *mVSG*-ES, and second the emergence of a single active *mVSG-*ES. Here, we present three independent lines of evidence to support this finding. First, in AnTat1.1E (EATRO1125) parasites from tsetse salivary glands, our inDrop data identified both metacyclic and pre-metacyclic cells, which allowed us to evaluate the expression of *mVSG* genes before monoallelic expression was established. Second, 10xGenomics data from Vigneron, O’Neill et al. (2020) using the RUMP503 strain were re-analyzed. Vigneron, O’Neill et al. (2020) concluded that each metacyclic cell expressed a single *mVSG,* as our re-analysis concluded. However, expression of multiple *mVSG* genes occurs before cells become metacyclic forms. The intermediate cell cluster identified in their study therefore likely represents the pre-metacyclic cells in our re-analysis. This analysis was only possible due to the addition of *de novo* assembled *mVSG* transcripts. Third, we used smRNA-FISH in an *in vitro* metacyclogenesis model using the Lister 427 strain. Taken together, our data from three strains analyzed by different techniques, strongly support our conclusions. This triple set of experimental evidence supports a dynamic process where many genes are initially transcribed at a low level, and as expression levels increase, a single gene emerges as the active *VSG*.

Metacyclic cells express a protein coat comprised of a single VSG type (Ramey-Butler, Ullu et al. 2015), suggesting that the multi-*<u>m</u>VSG* gene expression we observed is restricted to the transcript level in pre-metacyclic cells. Indeed, immunogold electron microscopy (Tetley, Turner et al. 1987) and immunofluorescence data (Rotureau, Subota et al. 2012) indicates that pre-metacyclic cells do not have VSG on their surface. According to the nomenclature from Tetley and Vickerman (1985), VSG protein appears on the surface of nascent metacyclic cells, a developmental stage between pre-metacyclic and metacyclic form cells. As no cell division occurs between pre-metacyclic/nascent/metacyclic differentiation, these data suggest that there is a temporal component to ensuring a single VSG coat which depends upon both the time required to establish a single active locus and the differentiation from pre-metacyclic to metacyclic cell. No data is available on the duration of the transition from pre-metacyclic to metacyclic cell, so it is not possible to estimate the rate at which a single *mVSG*-ES is chosen *in vivo*. Evidence from VSG switching experiments in bloodstream forms indicates that new VSG proteins are detectable on the surface within 12 h of a switch (Pinger, Chowdhury et al. 2017), suggesting that the establishment of monoallelic expression could occur within this time frame. During the transition from BSF to PCF following a tsetse bloodmeal, the VSG coat is removed and replaced with procyclin. The appearance of procyclin mRNA precedes the appearance of protein by approximately 6h (Roditi, Schwarz et al. 1989). If a similar delay exists with BARP to mVSG coat remodeling, this could potentially explain how parasites are able to express a single VSG type in their mVSG coats. Nevertheless, the exact parameters underlying these transitions remain to be determined.

In bloodstream form cells, a single *VSG* is transcribed from the active *VSG-*ES in the ESB (Navarro and Gull 2001). In metacyclic cells, the picture is less clear: RNA Pol I staining indicated that no ESB is detected at this stage. Metacyclic *VSG*-ES are much shorter than BSF *VSG*-ES (∼5 Kbp for *mVSG*-ES compared to ∼60 Kbp for BSF *VSG*-ES), therefore the absence of an ESB in metacyclic cells may be due to fewer RNA Pol I molecules at these loci (Ramey-Butler, Ullu et al. 2015). Inferring from the characteristics of the bloodstream ESB, we can speculate on nature of *VSG* gene expression regulation in metacyclic cells. The ESB is a bi-partite nuclear body consisting of the mRNA trans-splicing machinery and the actively transcribed *VSG*-ES (Faria, Luzak et al. 2021). VEX1 and VEX2 are markers of the ESB, and their association with the ESB are key to the maintenance of VSG monoallelic expression. Perturbation of either or both protein’s expression leads to dramatic failures in monoallelic VSG expression control, with silent VSG-ES being transcribed and multiple VSGs expressed on the surface (Glover, Hutchinson et al. 2016, Faria, Glover et al. 2019). Pharmacological inhibition of transcription disperses the VEX1 and VEX2 and RNA Pol I foci (Glover, Hutchinson et al. 2016, Kerry, Pegg et al. 2017, Faria, Glover et al. 2019, Faria, Luzak et al. 2021), suggesting that the ESB requires active transcription. These data lead to a picture of the ESB as a structure which emerges from, and is stabilized by transcription of a *VSG* gene.

We propose a model for the establishment of monoallelic expression in metacyclic cells (Figure 7) where the multi-*mVSG* expression profiles observed in this study correlate to an initial step in a “race” between each *mVSG-*ES (Figure 7A) to obtain a sufficient level of transcription (Figure 7B) which leads to the down regulation of other sites. We propose that following asymmetric division of attached epimastigote parasites, pre-metacyclic cells begin transcribing multiple *mVSG*-ES. This would likely in turn recruit VEX1 and VEX2 proteins to these loci, triggering a positive feedback loop, where VEX proteins recruit the mRNA splicing machinery, stabilize the ESB and drive further transcription. This could lead to suppression of transcription at other *mVSG*-ES by depletion of these limiting factors, or by the previously proposed “RNA silencing”, in which transcripts from the active *VSG* silence transcription from other *VSG-*ES (Glover, Hutchinson et al. 2016).

**Figure 7.**
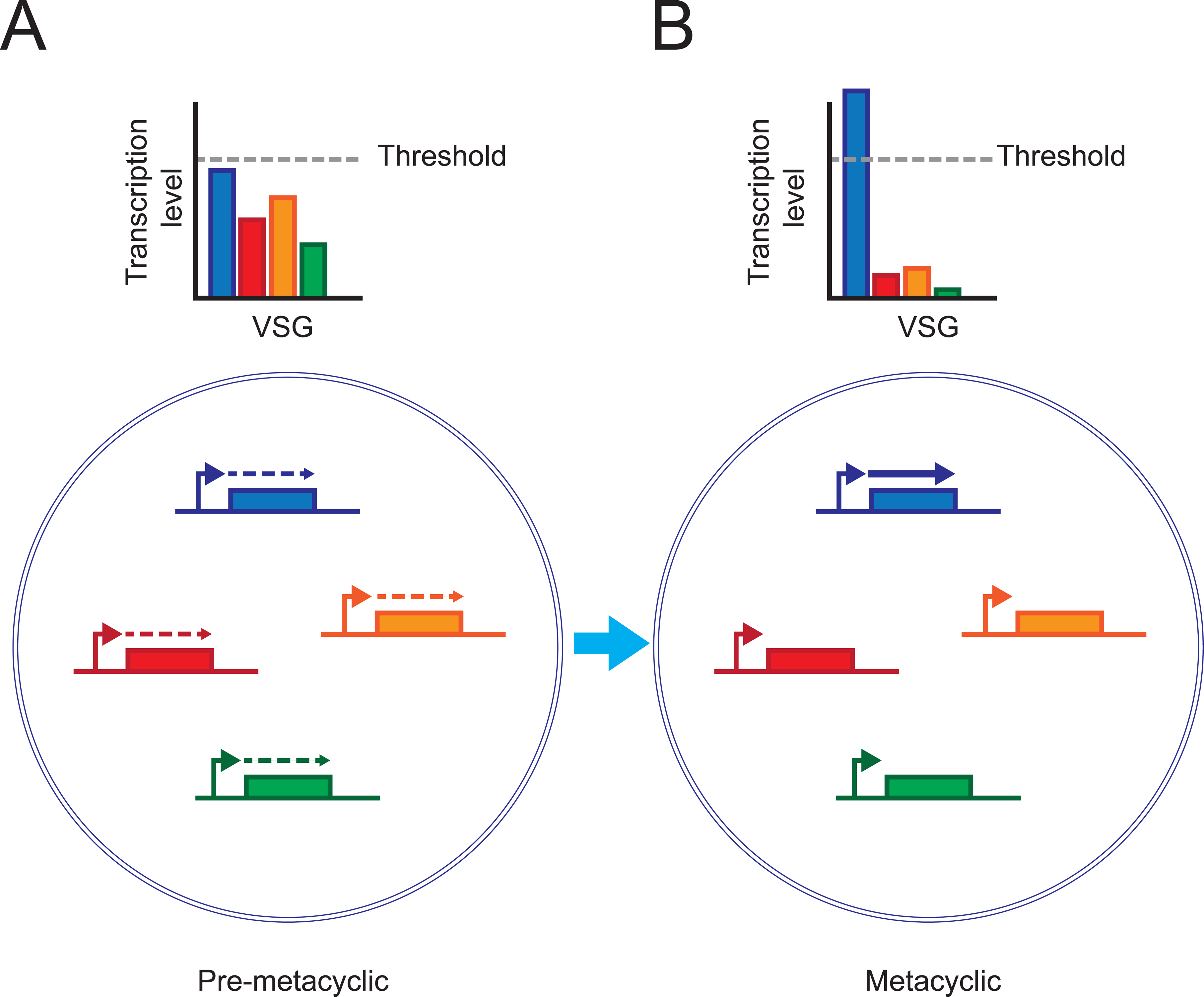
Model of the initiation of *VSG* monoallelic expression during metacyclogenesis. A. Pre-metacyclic cells initiate a transcriptional race for monoallelic expression where a single *VSG* gene must reach a threshold of transcriptional activity. B. In metacyclic cells, this threshold has been reached for a single VSG that is active, while the transcription of the other *mVSG*s is silenced.

We note a striking similarity between the dynamics controlling the establishment of monoallelic expression in both *mVSG* and olfactory neuron *OR* gene choice (Hanchate, Kondoh et al. 2015). In Metazoa, monoallelic expression underpins the expression of olfactory receptors (*ORs*). It is initiated during neuronal development in the olfactory bulb, and establishment of monoallelic expression is preceded by expression of multiple olfactory receptors in early mature olfactory neurons (Hanchate, Kondoh et al. 2015). A combination of DNA enhancer networks and an epigenetic trap, where the expression of an OR triggers a feedback loop which inhibits further activation after the first gene, establishing a single active OR. Here, during development of olfactory neurons, all *OR* genes are modified with heterochromatic histone trimethyl marks H3K4me3 and H4K20me3 (Magklara, Yen et al. 2011). The lysine demethylase LSD1 is then transiently expressed during development, activating a single OR (Lyons, Allen et al. 2013). A feedback signal triggered by OR expression utilizing the misfolded protein response inhibits further OR gene activation (Dalton, Lyons et al. 2013).

Both systems have intermediate developmental stages where transcripts from multiple *mVSG/OR* genes are detected, which then resolve to monoallelic states. Further, the establishment of monoallelic expression of *OR* genes is dependent on the structured and temporal control of histone methylation (Magklara, Yen et al. 2011). Similarly, in trypanosomes, the histone methyltransferase DOT1B is required for strict control of monoallelic expression following a transcriptional switch (Figueiredo, Janzen et al. 2008). In BSF cells, ectopic expression of a second VSG leads to silencing of the active VSG that is dependent on DOT1B (Batram, Jones et al. 2014). In the race model, the *mVSG-*ES who lost the “race” for transcription would be silenced in a DOT1B dependent manner following the initial race, consistent with the two-stage process observed in BSF cells. These similarities suggest common solutions to the problem of monoallelic expression which deserve further investigation. Through both experimental data, and re-analyses of published data, we are able to uncover the dynamics underlying the establishment of monoallelic VSG expression. Our data provide a landscape to frame these future investigations into the establishment of single antigen expression in trypanosomes.

## Materials and methods

### Tsetse fly infections and dissections

Tsetse flies (*Glossina morsitans morsitans*) were maintained at 27°C and 70% hygrometry in Roubaud cages, in groups of 50 male flies per cage. Teneral flies were infected with *Trypanosoma brucei* Antat1.1E (EATRO1125) during their first meal. Stumpy form trypanosomes were resuspended at 10^6^ cells per ml in SDM-79 (Brun and Schonenberger 1979) supplemented with 10% FBS (dominique Dutscher) and 60 mM N-acetylglucosamine (Peacock et al., 2006). Flies were allowed to feed on infected media through a silicone membrane. Following infection, flies were then maintained until dissection by feeding four times per week on sheep’s blood in heparin. Flies were dissected as described previously (Rotureau, Subota et al. 2012). All salivary glands were collected (uninfected and infected) in trypanosome dilution buffer on ice and midguts scored for the presence of trypanosomes. Salivary glands were manually broken using tweezers before being passed through a 70 µm cell strainer (ClearLine) to remove the salivary glands. Trypanosomes were counted on a hemocytometer and diluted to 9.4 x 10^4^ cells per ml. The final concentration was reached by adding 18 µL of OptiPrep™ Density Gradient Medium to 100 µL of cell suspension (final concentration 15 % vol/vol) immediately prior to encapsulation. Cells were maintained on ice throughout this process.

### Microscopy

Live trypanosomes in SDM-79 (Brun and Schonenberger 1979) were spread on glass slides and imaged on a DMR microscope (Leica) equipped with a EMCCD camera (C-9100, Hamamatsu). The camera was controlled using µManager and a plugin for Hamamatsu cameras (Edelstein, Tsuchida et al. 2014) and captured images were edited using Fiji is just ImageJ (FIJI) (Schindelin, Arganda-Carreras et al. 2012). For the viability assay, cells were scored for motility in a hemocytometer using an inverted phase contrast microscope.

### Cell Encapsulation

InDrop emulsions were prepared as described in Zilionis, Nainys et al. (2017). A detailed protocol compatible with the microfluidic designs described by Zilionis, Nainys et al. (2017) is available on (10.5281/zenodo.3974628). Monodisperse droplets were produced by injecting the 3 aqueous phases consisting of the cells mix, RT and lysis mix and barcoded beads mix, and a fluorinated oil phase containing fluorosurfactant into a microfluidic chip with controlled flow rates, creating an emulsion. Polydimethylsiloxane (PDMS) chips were fabricated as previously described (Mazutis, Gilbert et al. 2013) at the Institut Pierre-Gilles de Gennes (IPGG) in Paris. After plasma bonding, chips were made hydrophobic by manual injection of 2 % 1H,1H,2H,2H-perfluorodecyltrichlorosilane (Sigma-Aldrich) diluted in HFE 7500 (3M Novec).

Three aqueous phases, respectively for the cells, reverse transcriptase/lysis and BHMs (1CellBio) were prepared as follows; the cell mix contained 100 µL of cells diluted at a concentration of 9.4 x 10^4^ cells per ml in trypanosome dilution buffer (TDB) (5 mM KCl, 80 mM NaCl, 1 mM MgSO_4_, 20 mM Na_2_HPO_4_, 2 mM NaH_2_PO_4_, 20 mM glucose, pH 7.4) supplemented with 18 µL Optiprep (1CellBio). The RT mix contains 1.3 RT premix (1CellBio), 11 mM MgCl_2_ (1CellBio), 6.9 mM DTT (Invitrogen), 1.13 U/µL SUPERase IN RNAse Inhibitor (Life Technologies), 20.7 U/µL Superscript III (SSIII) (Invitrogen) and 1 µM DY-647 (Dyomics), the beads mix contains BHMs (1CellBio) packed in 1x Gel Concentration Buffer (1CellBio). The commercial BHMs carry photocleavable barcoded primers, containing, in order, a T7 promoter sequence for IVT, an Illumina adaptor, a cellular barcode, a 6 nt UMI and a 18TVN primer site, as described (Zilionis, Nainys et al. 2017). The closely packing of the BHBs allows 80-90% loading of a single bead per droplet (Abate, Chen et al. 2009).

The 1 ml syringes (Omnifix) for the cells, reverse transcriptase/lysis and BHMs were loaded with mineral oil (Sigma-Aldrich) and connected to a 200 µL pipette tip prefilled with mineral oil using ∼30 cm of polytetrafluoroethylene (PTFE) 0.56 mm tubing and a PDMS plug connecting the tubing to the tip. Mineral oil was then ejected from the syringe until it entirely filled the pipette tip prior to aspiration of each aqueous phase. Each reagent was aspirated on ice into the 200 µL pipette tips filled with mineral oil, just before encapsulation. Cells were maintained on ice throughout the procedure by placing the pipette tip holding the cells in ice. A short length (15 cm) of PFTE 0.56 mm tubing then connected this tube to the microfluidic chip. The RT/lysis and BHB pipette tips were directly inserted into the PDMS chip, whereas the tubing carrying cells from the pipette on ice was inserted into the PDMS chip. The syringe for the oil phase was filled with carrier oil (HFE-7500 fluorinated oil (3M Novec) containing 2% w/w 008-FluoroSurfactant (RAN Biotechnologies)) and connected to the PDMS chip by PFTE 0.56 mm tubing.

Droplet production and beads loading were followed using an inverted microscope (Nikon Eclipse Ti) and High-Speed Camera (Phantom). Flow rates for droplet production were controlled with syringe pumps (neMESYS 290N Low pressure Syringe Pump, Cetoni). Droplets size and frequency of production were monitored using laser optics excitation and detection and a soluble fluorophore DY-647 present in the droplets (Dyomics) as described (Mazutis, Gilbert et al. 2013). Flow rates of 250 µL/h, 250 µL/h, 60-80 µL/h and 400-500 µL/h for the cells, RT/lysis and BHMs and oil, respectively were maintained. A 5 cm PTFE tubing (0.3 mm inner diameter) connects the chip outlet to a 1.5 mL collection tube, prefilled with 300 µL of mineral oil. Emulsions were collected during 15 minutes on ice, corresponding to approximately 4,000 drops containing both a cell and and a BHM.

### Generation and sequencing of barcoded cDNA libraries

We made two minor alterations to the molecular biology workflow described by Zilionis, Nainys et al. (2017): Our protocol omits the RNA fragmentation step following IVT of double stranded cDNA products, and we modified the primer used to reverse transcribe amplified RNA (aRNA) to make our libraries compatible with standard Illumina sequencing primers. These alterations are unlikely to affect the overall performance of inDrop, however.

#### 1. Recovery of first strand cDNA

Barcoded cDNA libraries were prepared according to the protocol by Zilionis, Nainys et al. (2017) with small exceptions. Following encapsulation and ultra violet primer release, emulsions were incubated at 50°C for 2h and then 70°C for 15 minutes. Following reverse transcription (RT), emulsions were partitioned into approximately 2,000 cells per partition before breaking the emulsion by adding one drop (3µL) of 1H,1H,2H,2H-Perfluorooctanol (Sigma-Aldrich) (Zilionis, Nainys et al. 2017). cDNA was then stored at −80°C until further processed.

Post-RT material was thawed on ice and 20µL of dH_2_O (nuclease free) was added and the solution centrifuged at 16,000 g for 5 minutes at 4°C. The aqueous phase was then applied to a 0.45 µm filter spin column (Corning CoStar Spin-X, Sigma-Aldrich) to remove the BHMs. The emulsion tube was then washed with 20µL dH_2_O to collect as much cDNA as possible and applied to the spin column. The column was centrifuged at 16,000 g for 5 minutes at 4°C. The emulsion tube was washed once more and supernatant collected by centrifugation once more.

#### 2. Removal of unused primers

The volume of cDNA was estimated and digested by adding 100 µL of digestion mix (79 µL H_2_O, 9 µL 10x Fast Digest Buffer (Thermo Scientific), 5 µL ExoI (20 U/µL, NEB), 7 µL HinFI (10U/µL, Thermo Scientific)) per 70 µL of cDNA and digested at 37°C for 30 minutes. Samples were then purified by adding 1.2x volume of room temperature AMPure XP beads (Beckman Coulter), incubating 5 minutes at room temperature, collecting the beads on a magnet and then 3 washes with freshly prepared 80% ethanol (room temperature), followed by air drying, according to the manufacturer’s instructions. cDNA was then eluted in 17 µL of dH_2_O.

#### 3. Second strand synthesis

Second strand synthesis (SSS) was performed using the NEBNext Ultra II Non-Directional RNA Second Strand Synthesis Module (E6111S). 2 µL of SSS buffer and 1 µL of SSS enzyme mix was added to 17 µL of cDNA and the reaction was incubated at 16°C for 2h30m in a PCR machine with the lid set to 16°C. cDNA was purified again using 1.2x AMPure XP beads (as above) and eluted in 7µL of DNA elution buffer (Zilionis, Nainys et al. 2017).

#### 4. *In vitro* transcription and reverse transcription of amplified RNA (aRNA)

Double stranded cDNA was then amplified using the HiScribe T7 Quick High Yield RNA Synthesis Kit (E2050S). To the 8µL of double stranded cDNA, 10 µL of 10x NTPs buffer mix, 2 µL of T7 enzyme and 1 µL of SUPERase IN RNAse Inibitor (20 U/µL). were added and the reaction incubated at 37°C with the lid set to 50°C for 14-16 hours. The size and concentration of amplified RNA was verified using a BioAnalyzer RNA pico chip (Agilent) using the mRNA pico program. Approximately 5 ng of aRNA was then reverse transcribed by adding 1 µL of 10 mM dNTPs, 2 µL of Rd2N6 (see primer list) and dH_2_O to 13 µL. The RT reaction was heated to 70°C for 3 minutes and then put on ice for 1 minute. Finally, 4 µL of 5x first strand buffer, 1 µL of 0.1M DTT, 1 µL of SSIII (200 U/µL) and 1 µL of SUPERase IN RNase inhibitor (20U/µL). RT reactions were incubated for 5 minutes at 25°C, followed by 1h at 50°C and heat inactivated at 70°C for 15 minutes. cDNA was purified by adding 20 µL of dH_2_O then purified with 1.2 x AMPure XP beads and eluted with 20 µL dH_2_O. 10 µL of this was then stored at −80°C as a pre-PCR backup.

#### 5. Library PCR

The remaining 10 µL of cDNA was used for the library PCR. Libraries were PCR amplified using KAPA HiFi HotStart ReadyMix (Kapa Biosystem, KK2601). 25 µL of PCR mix containing 12,5 µL of 2x KAPA HiFi Master mix, 10 µL of cDNA, together with 2.5 µL of 5 µM P5-Rd1 and P7-Rd2 primers (final concentration of 500 nM). Library indices were in the P7-Rd2 primer. Primer sequences are available in Supplementary table 4. The libraries were amplified using the following protocol; initial denaturation 5 min at 95°C, then 2 cycles of denaturation 98°C / 20 s, annealing 55°C / 30s and extension 72°C / 40 s, followed by 10 cycles of denaturation 98°C / 20 s, annealing 65°C / 30 s and extension 72°C / 40 s and final extension 5 min at 72°C. They were then size selected using 0.7x volume of AMPure XP beads at room temperature prior to quantification and sequencing.

#### 6. Library quantification and sequencing

Libraries were quantified using the KAPA library quantification kit for Illumina (Kapa Biosystem, KK4824) according to the manufacturers’ protocol. Bulk libraries (Figure 1B) were sequenced on the Illumina MiSeq platform with a MiSeq V3 150 cycle cartridge at the Institut Curie. Single cell barcoding libraries were sequenced on the Illumina NextSeq platform at the Institut Pasteur using a Mid-output 150 cycle cartridge. Sequencing of single cell libraries at BGI was performed on the MGIseq-2000 following conversion to DNBseq libraries (Senabouth, Andersen et al. 2020). Details of sequencing outputs including read lengths and number of reads generated can be found in Supplementary table 5.

### Trypanosoma brucei transcriptome

We assembled a non-redundant transcriptome from the T. brucei TREU927 reference strain (Berriman, Ghedin et al. 2005). As only 60% of the trypanosome transcriptome has annotated 3’ UTRs, we decided to extend each gene’s annotated 3’ extremity to the 5’ extremity of the neighboring gene. This was done by a combination of scripted extension using the following rules. 1. If the proceeding gene is on the same strand, and within 2.5 kbp, the gene is extended to the base before the proceeding gene. 2. If the proceeding gene is over 2.5 kbp then the gene is extended to 2.5 kbp downstream. 3. If the gene already extended into the proceeding gene, then no change was made. 4. Similarly if the proceeding gene is on the opposite strand, no extension was made. Following extension of the entire gene list, extended transcripts were filtered using a non-redundant gene list (Alsford, Turner et al. 2011).

To obtain sequences of metacyclic *VSG* genes expressed in the salivary glands, we aligned our data to the VSGnome (including flanks) of the Antat1.1E (EATRO1125) strain made available by Cross (2017) here (http://129.85.245.250/index.html), using bowtie2 (Langmead and Salzberg 2012) and selected the top 8 gene most abundant transcripts. These are Tbb1125VSG-1401 (Genbank: KX699657), Tbb1125VSG-1476 (Genbank: KX699716), Tbb1125VSG-1654 (Genbank: KX699860), Tbb1125VSG-385 (Genbank: KX699226), Tbb1125VSG-393 (Genbank: KX698711), Tbb1125VSG-4564 (Genbank: KX700928), Tbb1125VSG-4862 (Genbank: KX701110), Tbb1125VSG-4959 (Genbank: KX701187).

To this non-redundant transcriptome, we appended both a human transcriptome and the *G. morsitans morsitans* genome sequence (International Glossina Genome 2014). Our inDrop library preparations included 5% human B-cells as a positive control for the inDrop pipeline, and our bowtie2 index therefore contained the ENSEMBL release 85 human transcriptome (Yates, Achuthan et al. 2020). The alignment index was constructed with bowtie2-build (Langmead and Salzberg 2012). These files are available at 10.5281/zenodo.3974628.

### Generation and analysis of bulk inDrop data

Cultured procyclic form Antat1.1E (EATRO1125) trypanosomes were encapsulated using a standard inDrop microfluidic chip, except we replaced BHMs with a single primer, T7Rd2_linkerPolyTvN (Supplementary table 4). Libraries were generated as described by Zilionis, Nainys et al. (2017), and as described for single cell BHMs in this manuscript, except we excluded stages relating to filtration of BHMs. Libraries were sequenced on a MiSeq at the Institut Curie (Paris) next generation sequencing platform.

Bulk data were aligned to the trypanosome genome sequence using bowtie2 (Langmead and Salzberg 2012), and resulting bam files manipulated using samtools (Li, Handsaker et al. 2009). Metagene plots and heat maps were generated using deeptools2 (Ramirez, Ryan et al. 2016). Gene coordinates used in the metaplot were the 5’and 3’splicing and polyadenylation signal sites (or start and stop codons if unannotated for a particular gene). Bin size was set to 10 bp and the 500 bp region downstream from each gene are unscaled. The gene list was filtered to a non-redundant set (Alsford, Turner et al. 2011).

### inDrop pre-processing pipeline

To permit more usability with bespoke genome sequences (no reference sequence is available for the strain used here for tsetse infections), we generated a new inDrop data analysis pipeline for the analysis of trypanosomes, using bowtie2 and UMI-tools (Langmead and Salzberg 2012, Smith, Heger et al. 2017), rather than bowtie and the inDrop analysis pipeline (Klein, Mazutis et al. 2015). Pre-processing of inDrop data was performed using the UMI-tools package (Smith, Heger et al. 2017). A bash script is available at (10.5281/zenodo.3974628) which performs all steps of the pre-processing. The script executes 5 steps: 1. whitelist, 2. fastqsplitter, 3. extract, 4. align and 5. count. Whitelist generates a list of acceptable barcodes, using a regular expression to pattern match the cell barcode flanked by the common “W1” sequence (Klein, Mazutis et al. 2015, Zilionis, Nainys et al. 2017) permitting upto 6 errors in the W1, and requiring a string of 3 “T” nucleotides after the UMI. We set the error correction threshold for barcodes at 2, and set the expected number of cells at 2,000. We subset 10^6^ reads to perform the whitelist step (these same filtering parameters are maintained through our workflow). Fastqsplitter breaks the read files into a user-defined number of equal parts to allow downstream parallel processing. Extract then places the whitelisted cell barcodes from read 1 into the header of mRNA sequences in read 2, using the regex pattern matching parameters from whitelist. The script can initiate multiple instances of extract to reduce the time required for this stage. Align uses bowtie2 (Langmead and Salzberg 2012) to align the reads to the hybrid trypanosome, human transcriptomes and tsetse fly genome. Reads aligning to the tsetse genome were filtered out using samtools view (Li, Handsaker et al. 2009). Finally count generates count matrices at mapping quality 0 and 40. Higher mapping stringency was used for *VSG* counts to ensure accurate mapping.

### Analysis of single cell RNA-seq data

Single cell count matrices were analysed using the Seurat R package (Butler, Hoffman et al. 2018, Stuart, Butler et al. 2019). Data were normalized using the single cell transform algorithm (Hafemeister and Satija 2019). Complete R scripts are available in (10.5281/zenodo.3974628) to generate all the figures in the manuscript. Count tables from mapq 0 and mapq 40 matrices were merged into a single matrix by removing VSG counts with mapq 0 and merging the mapq 40 counts to this reduced matrix. Valid barcodes were filtered by a minimum gene cut-off of 300. VSG UMI count data were extracted using fetchData, accessing the “counts” slot of the RNA assay.

To assess the sequencing depth, we employed data subsetting, as done previously by Zhang, Li et al. (2019). Data sets were subset using samtools and UMI totals per barcode extracted in R, retaining only valid barcodes (R script on 10.5281/zenodo.3974628). To calculate the number of mapped reads per cell, a custom script was developed to count the number of aligned reads per barcode (available on 10.5281/zenodo.3974628). Subset reads were then calculated as a fraction of the total. To calculate the Levenshtein distance between valid barcodes, the barcodes for replicate 1 were extracted and all possible pairs were evaluated using a python script (10.5281/zenodo.3974628). Histogram and plotting were performed in R. To estimate the expected rate of doublets a Poisson series was generated corresponding to the predicted occupancy of cells in droplets. This was then compared to the frequency of multi-*VSG* expression observed in the pre-metacyclic cell cluster. The metabolism analysis was performed using metabolic funtion annotations and schematic for glycolysis and TCA cycle from trypanocyc (Shameer, Logan-Klumpler et al. 2015). Scaled (Z-scores) are from the SCT assay of Seurat in the “scaled.data” slot.

### Single molecule RNA-FISH

Overexpression of RBP6 was performed using PT1^RBP6-OE^ cells (a kind gift provided by Lucy Glover, Institut Pasteur). PT1 cells are equivalent to the 2T1 BSF (Alsford, Kawahara et al. 2005). Wild type Lister 427 PCF cells were transfected with pHD1313 (Alibu, Storm et al. 2005). Second, a ribosomal spacer was targeted with pRPH, integrating a hygromycin resistance cassette and eGFP reporter gene (GFP tagged Sir2.3) with a RNA Pol-I promoter under the control of a tet operator (Alsford, Kawahara et al. 2005). The RPH locus was then disrupted with pH3E (Alsford, Kawahara et al. 2005), replacing hygromycin with a puromycin resistance cassette, resulting in PT1 cells. Cells were induced by addition of 10 µg/ml of tetracycline to the culture medium and diluting every 24h to 2 x 10^6^ cells per ml. We made staggered inductions to obtain 3, 4 and 5 day induced cultures on the same day. We used the ViewRNA™ Cell Plus Assay Kit (Catalog number 88-19000-99, Thermo Fisher) for smRNA-FISH. The company designed probes based on the *VSG-397* and *VSG-531* or alpha tubulin coding sequences. We followed the manufacturer’s instructions with a few minor exceptions. Before settling cells onto slides, we chilled cultures on ice and then fixed in ice cold 1 % paraformaldehyde. Fixed cells were immediately washed in ice cold PBS by spinning 3x at 1,000 g for 10 minutes. Cells were settled onto 8-well glass chamber slides (Nunc™ Lab-Tek™ II Chamber Slide™ System) and stained according to the manufacturer’s protocol. A minimum of 100 µL was used to avoid drying samples during incubations. Cells were imaged on a widefield Zeiss Axio Imager.Z2 upright microscope equipped with an LED light source, and an Axiocam 506 camera. Images were analysed using FIJI (Schindelin, Arganda-Carreras et al. 2012, Rueden, Schindelin et al. 2017).

## Acknowledgements

We thank George Cross for making the Antat1.1E (EATRO1125) VSGnome available to the research community. Artur Scherf and the Biomics platform at the Institut Pasteur provided access to the Illumina NextSeq platform. Jessica Bryant and Sebastian Baumgarten provided valuable assistance with Illumina sequencing, as well as many useful discussions, and critical reading of the manuscript. Sophie Créno from the Institut Pasteur high performance computing facility helped with computational workflows. Christelle Travaillé aided with tsetse fly infections. We thank David Horn (University of Dundee) for valuable advice and discussions. We thank Lucy Glover, Serge Bonnefoy, and Aline Araujo Alves for critical reading of the manuscript. We are indebted to Eliane Thion and Lucy Glover (Institut Pasteur, Paris) for the RBP6 strain, to Alena Zíková (Institute of Parasitology, České Budějovice) for advice on inductions, and to Susanne Kramer (Universität Würzburg, Würzburg) for advice on smRNA-FISH.

## Data Availability

All sequencing data generated in this study is available from the European nucleotide archive, accession number PRJEB43345. All scripts and code used to generate the figures are available from Zenodo 10.5281/zenodo.3974628.

## Supplementary figure legends

**Figure 1-figure supplement 1.**
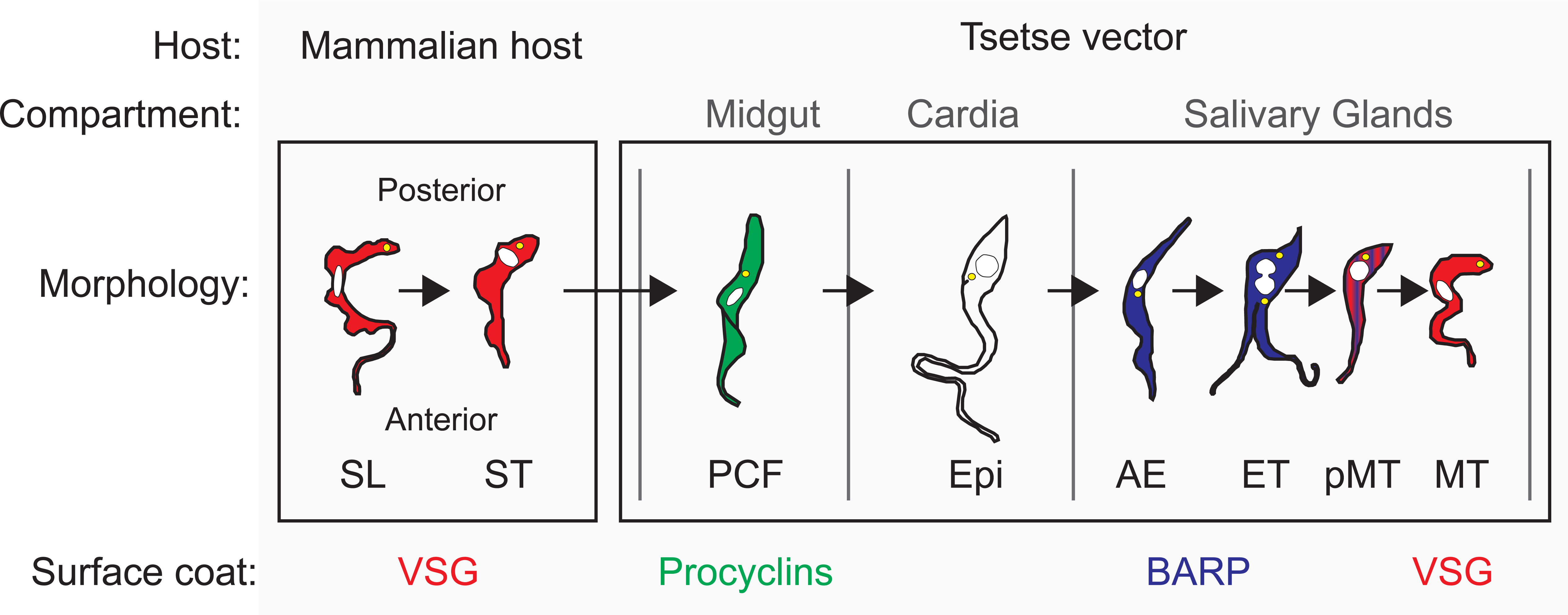
Schematic of the life cycle of *T. brucei*, adapted from Rotureau and Van Den Abbeele (2013). The morphology and surface protein expression of trypanosomes during differentiation is depicted in chronological order from left to right. Cell types shown here are SL: slender bloodstream form; ST: stumpy; PCF: procyclic form; Epi: epimastigote; AE: attached epimastigote; ET: epimastigote-trypomastigote dividing cell; pMT: pre-metacyclic cell; MT: metacyclic. The nucleus is depicted as a white circle / oval and the kinetoplast (mitochondrial DNA) as a yellow circle. In trypomastigote cells, the kinetoplast is posterior to the nucleus, e.g. in PCF. In epimastigote cells, the kinetoplast is anterior to the nucleus, e.g. in Epi. Cells are coloured according to the family of surface protein expressed.

**Figure 1-figure supplement 2.**
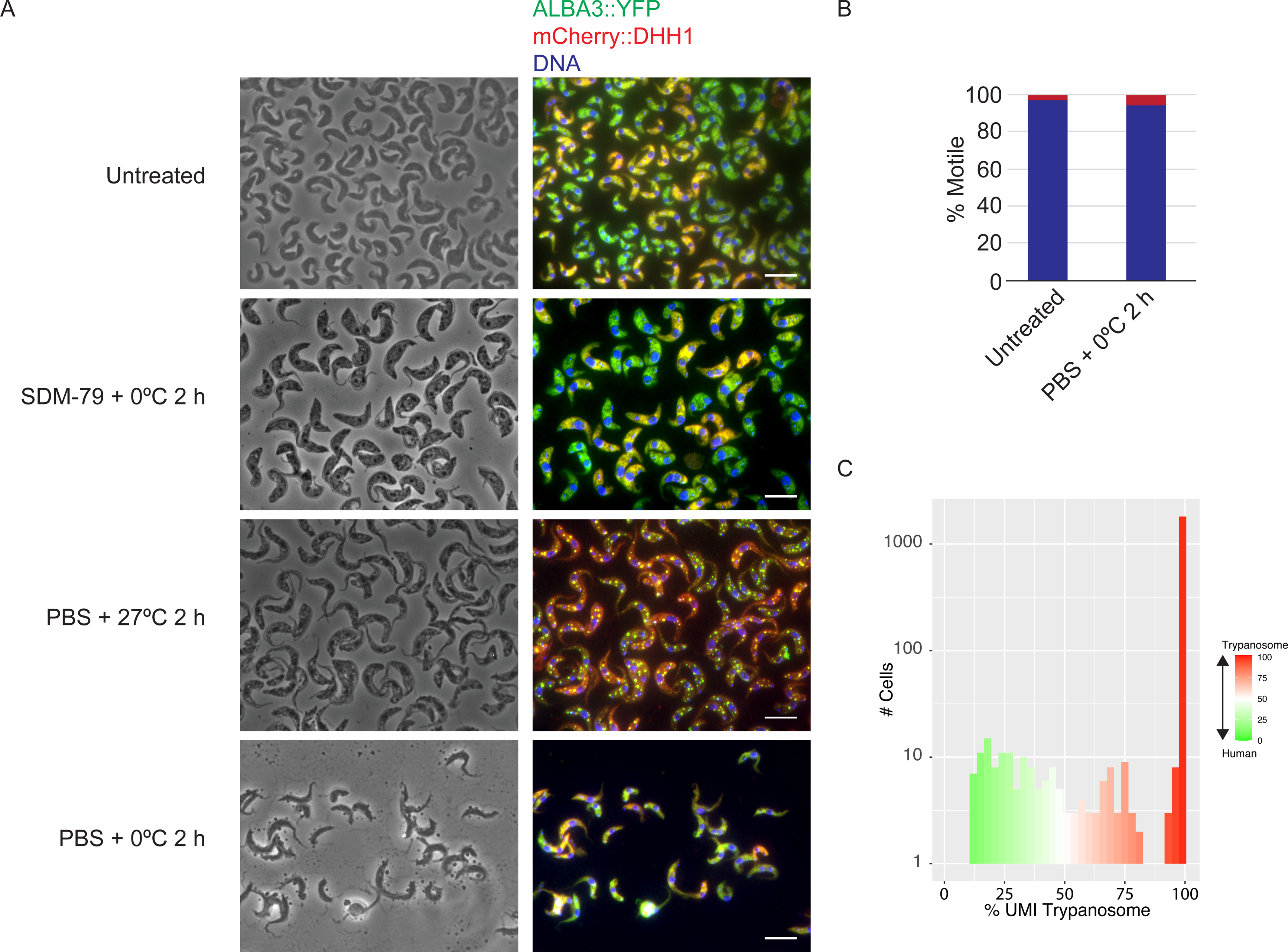
Survival and RNA metabolism of trypanosomes expressing the indicated fusion proteins in conditions mimicking the inDrop procedure. A. Trypanosomes collected by centrifugation and resuspended in either SDM-79 medium, or PBS and incubated at either 0°C or 27°C. Scale bar 10 µm. B. Results of live/dead assay based on trypanosome motility, by scoring live cells subjected to phase contrast microscopy. We assessed the ability of the parasites to survive starvation at 0°C for 2h in PBS. C. Histogram shows the percentage of stringently aligned reads (mapQ > 40) to the trypanosome or human transcriptomes.

**Figure 2-figure supplement 1.**
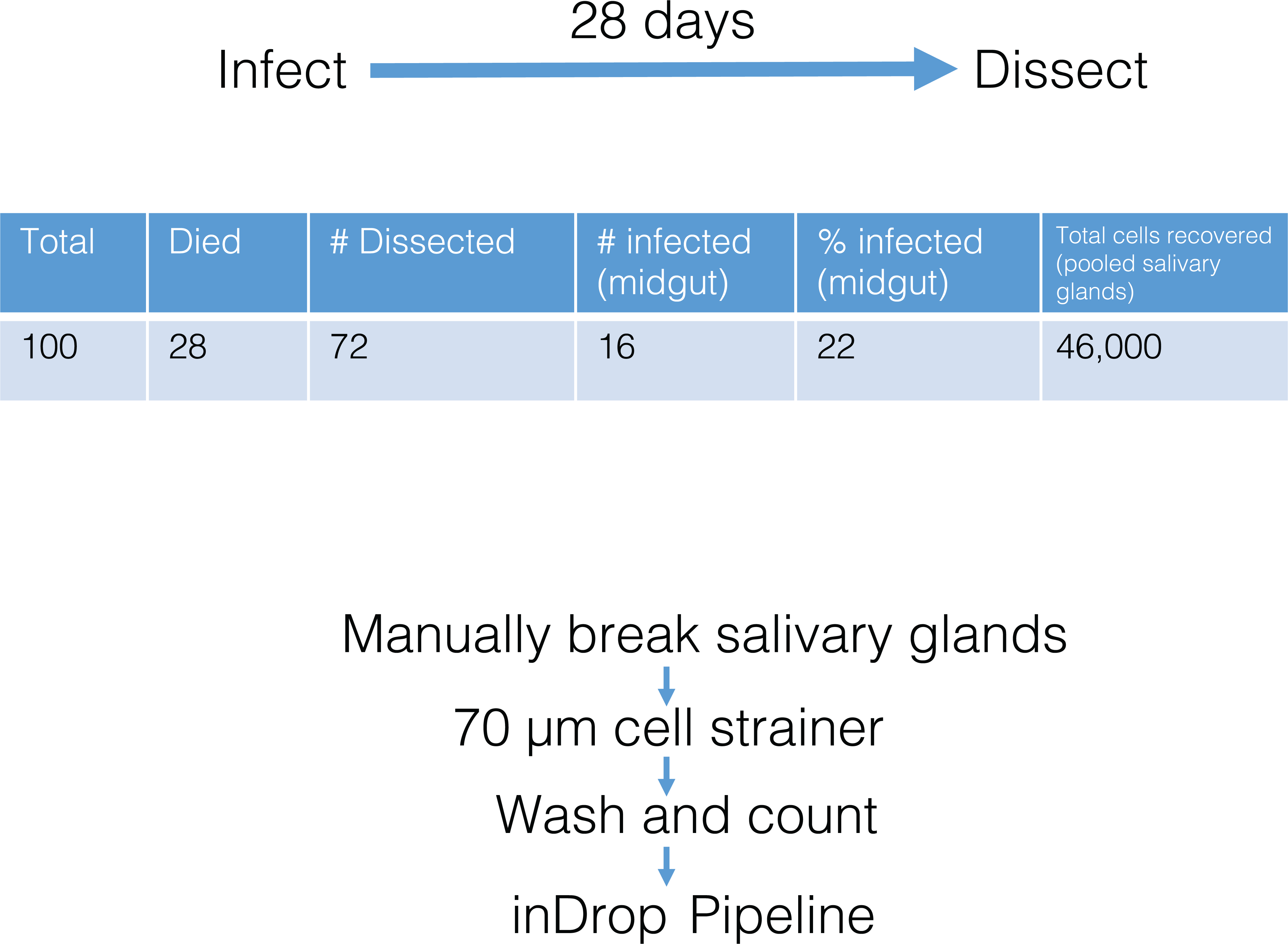
Details of tsetse fly infections. Flies were infected for 28 days. The table shows the characteristics of the dissection. Below is the workflow for dissections and sample collection.

**Figure 2-figure supplement 2.**
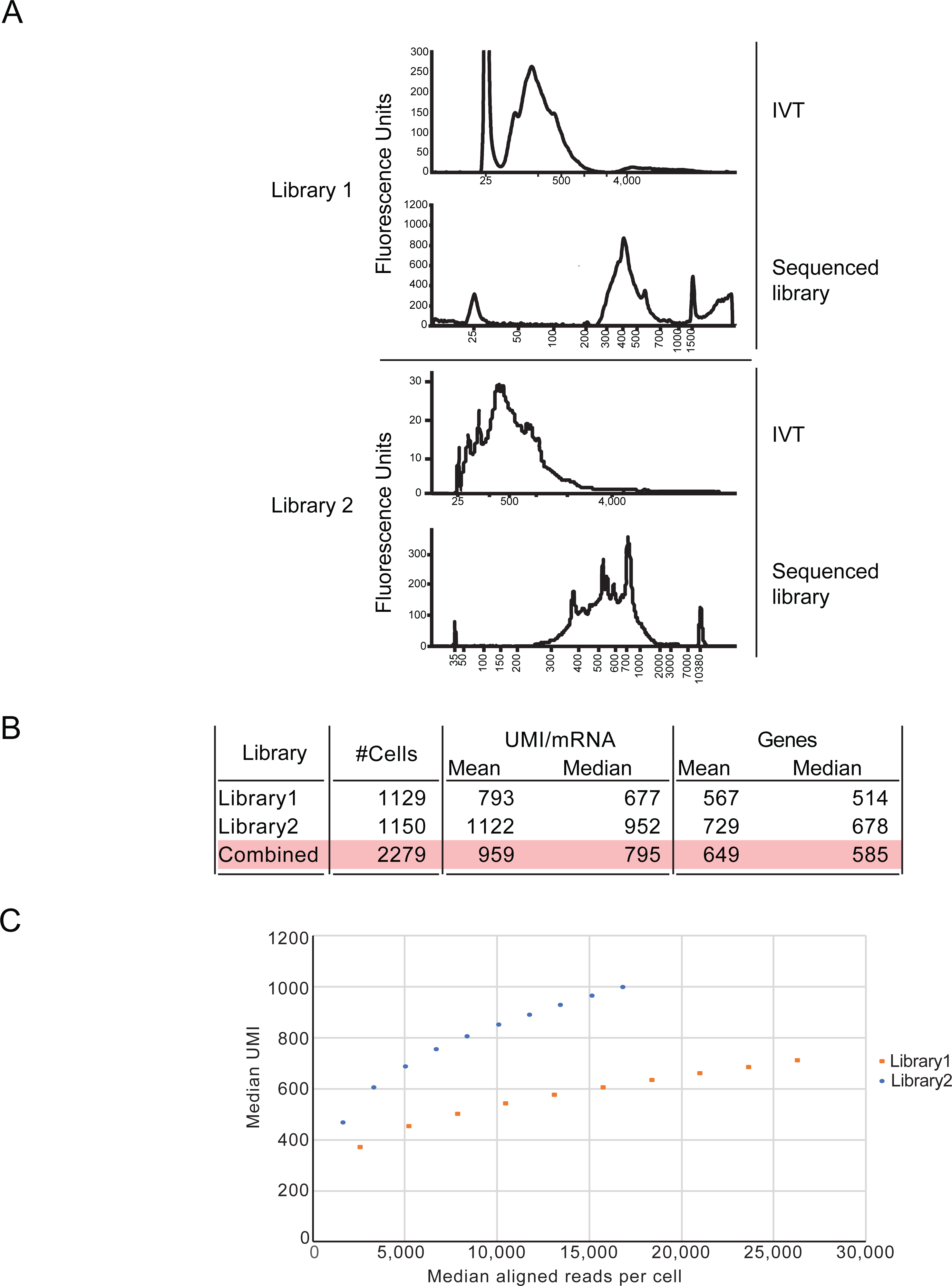
Library construction, basic statistics and subsetting analysis. A. Tapestation or Bioanalyzer traces for IVT and library PCR reactions to construct libraries sequenced in this study. B. Table shows the number of cells recovered, and key parameters including UMIs and genes captured. C. Subsetting analysis of the data. Reads were subsampled using samtools (Li, Handsaker et al. 2009) in 10% intervals from 10% to 100% before being re-counted using UMI-tools (Smith, Heger et al. 2017). Aligned reads per cell were counted using a custom script (10.5281/zenodo.3974628).

**Figure 2-figure supplement 3.**
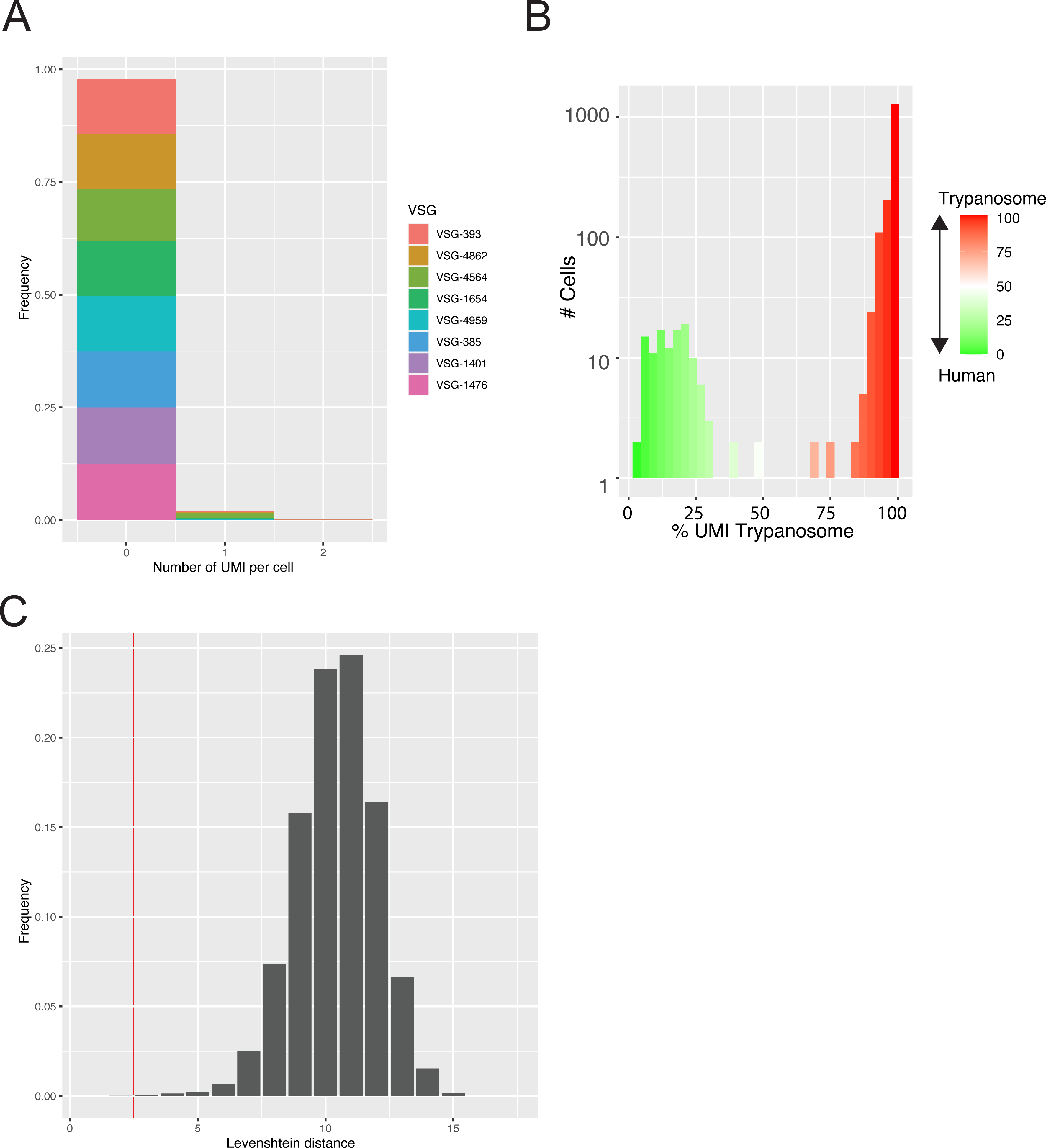
Analysis of possible technical artefacts. A. Ambient RNA analysis. Plot shows the frequency of *VSG* UMI detection in a cluster of Ramos cells. B. Histogram shows the percentage of stringently aligned reads (mapQ > 40) to the trypanosome or human transcriptomes. C. Barcode switching analysis. The plot shows the frequency of Levenshtein distances (number of changes required to permute one barcode to another) between all pairs of barcodes in replicate 1 library 1 (1,129 barcodes). Red line shows cut-off for error correction.

**Figure 3-figure supplement 1.**
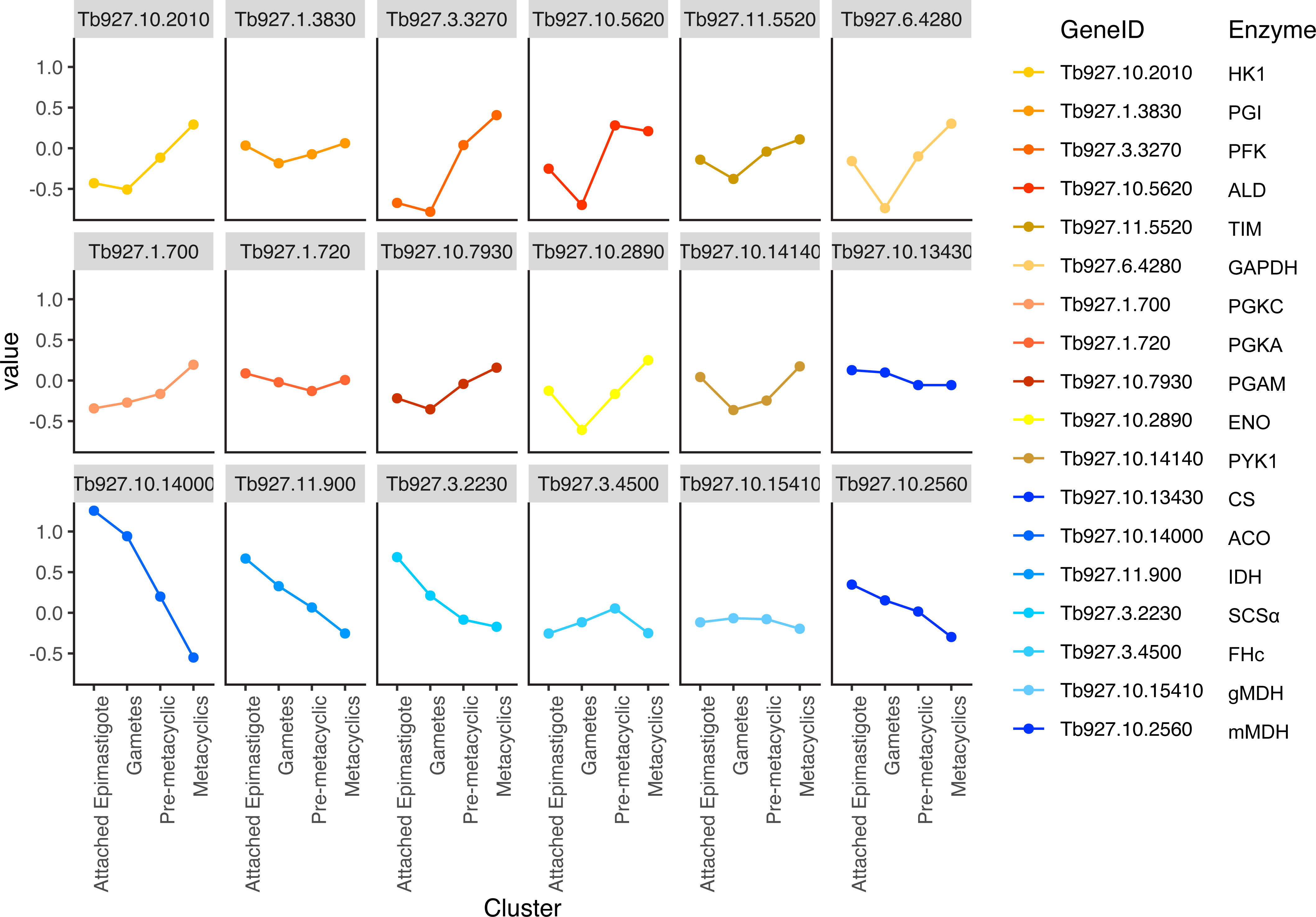
Analysis of metabolic enzymes in salivary gland clusters. As in Figure 3B except each gene is plotted individually.

**Figure 5-figure supplement 1.**
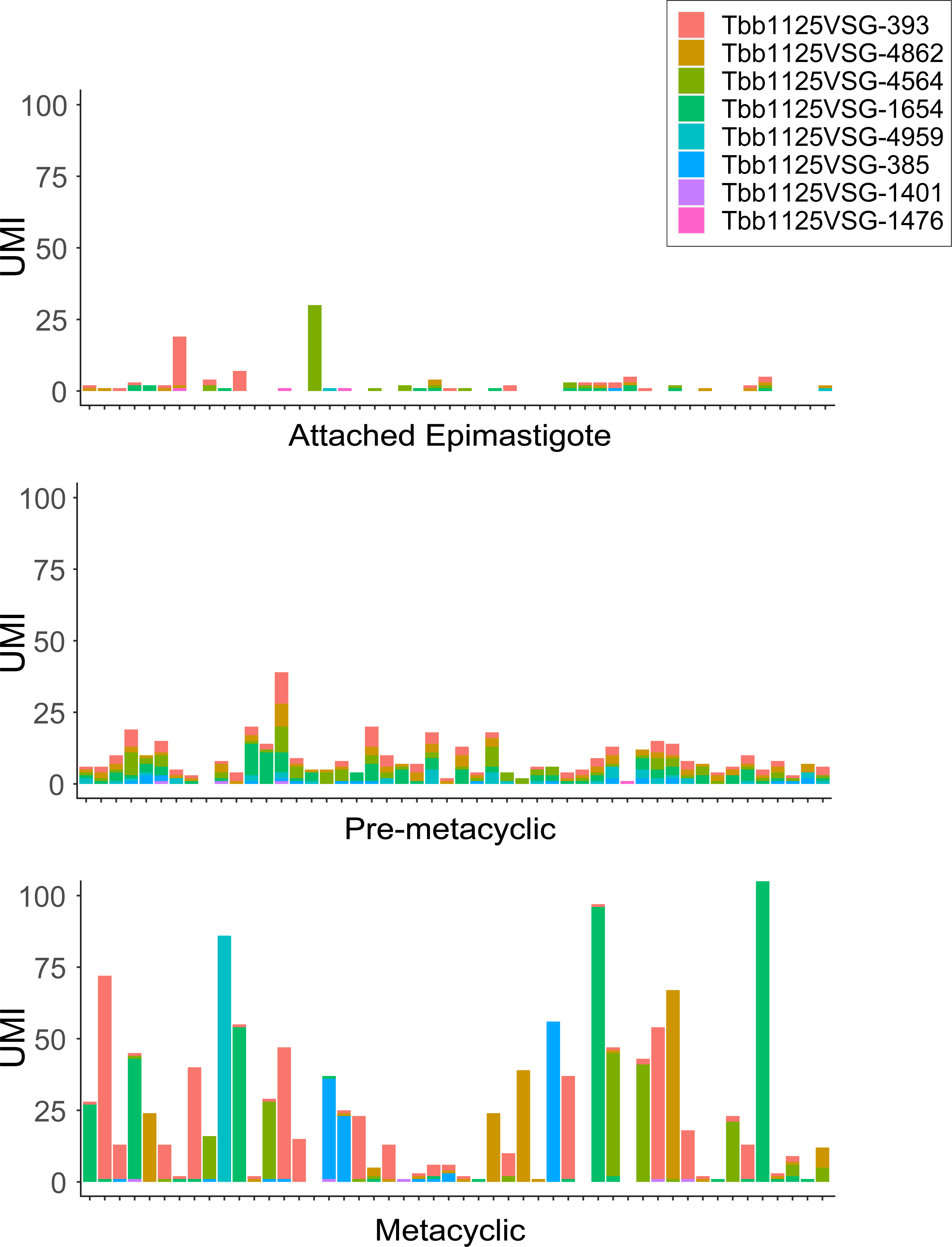
VSG expression analysis for attached epimastigote, pre metacyclic and metacyclic clusters. Data presented have not been filtered (no minimum UMI cutoff per cell) and consist of 50 random cells. Units are raw (non-normalised and unscaled) UMI counts per cell. All VSG counts use a stringent mapping quality (MapQ40).

**Figure 5-figure supplement 2.**
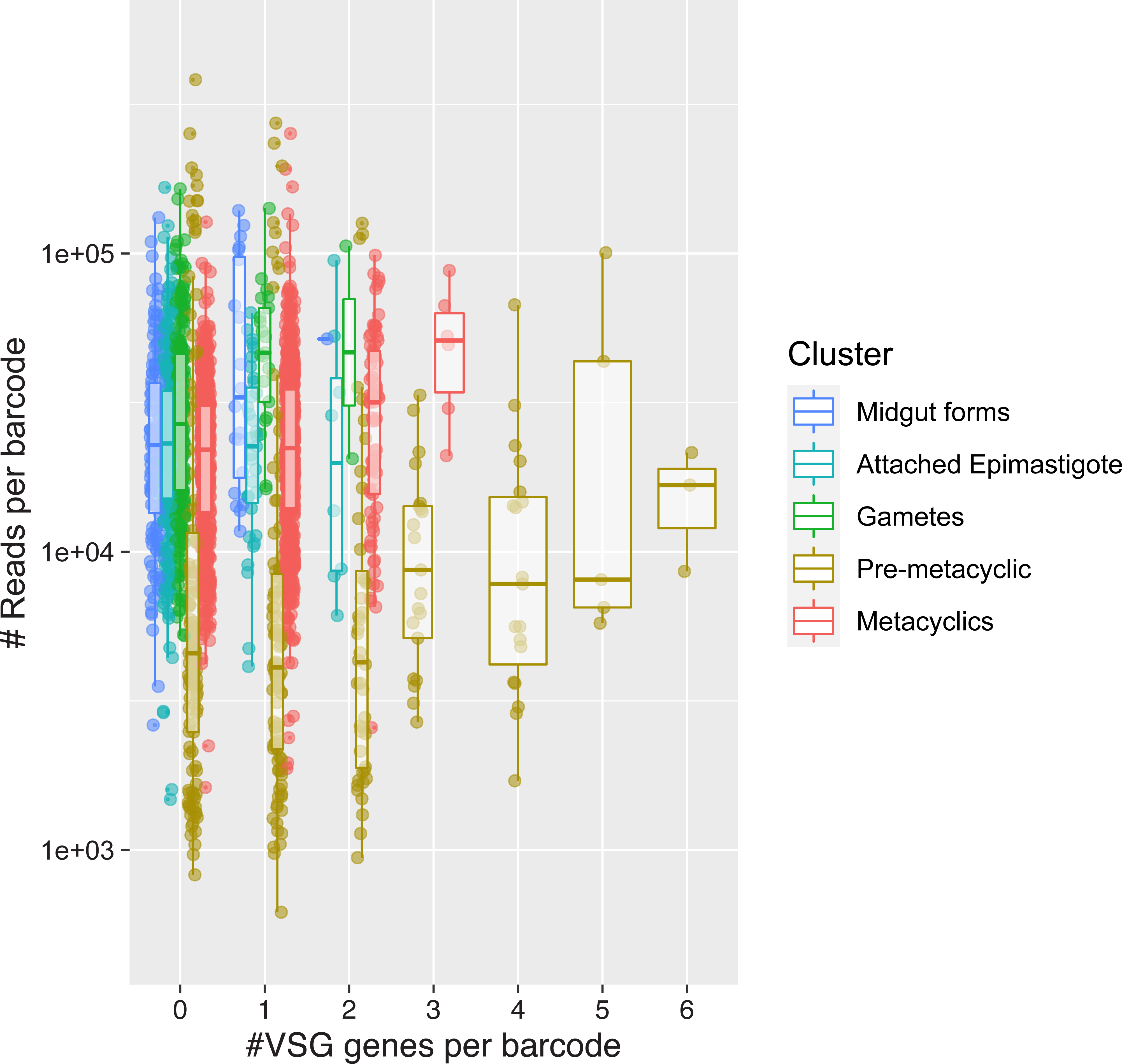
Plot shows the number of *VSG* genes detected per barcode versus the aligned reads per barcode stratified by cluster. Boxplots show bounds are 25^th^ 50^th^ and 75^th^ percentiles. Individual barcodes are plotted as dots.

**Figure 5-figure supplement 3.**
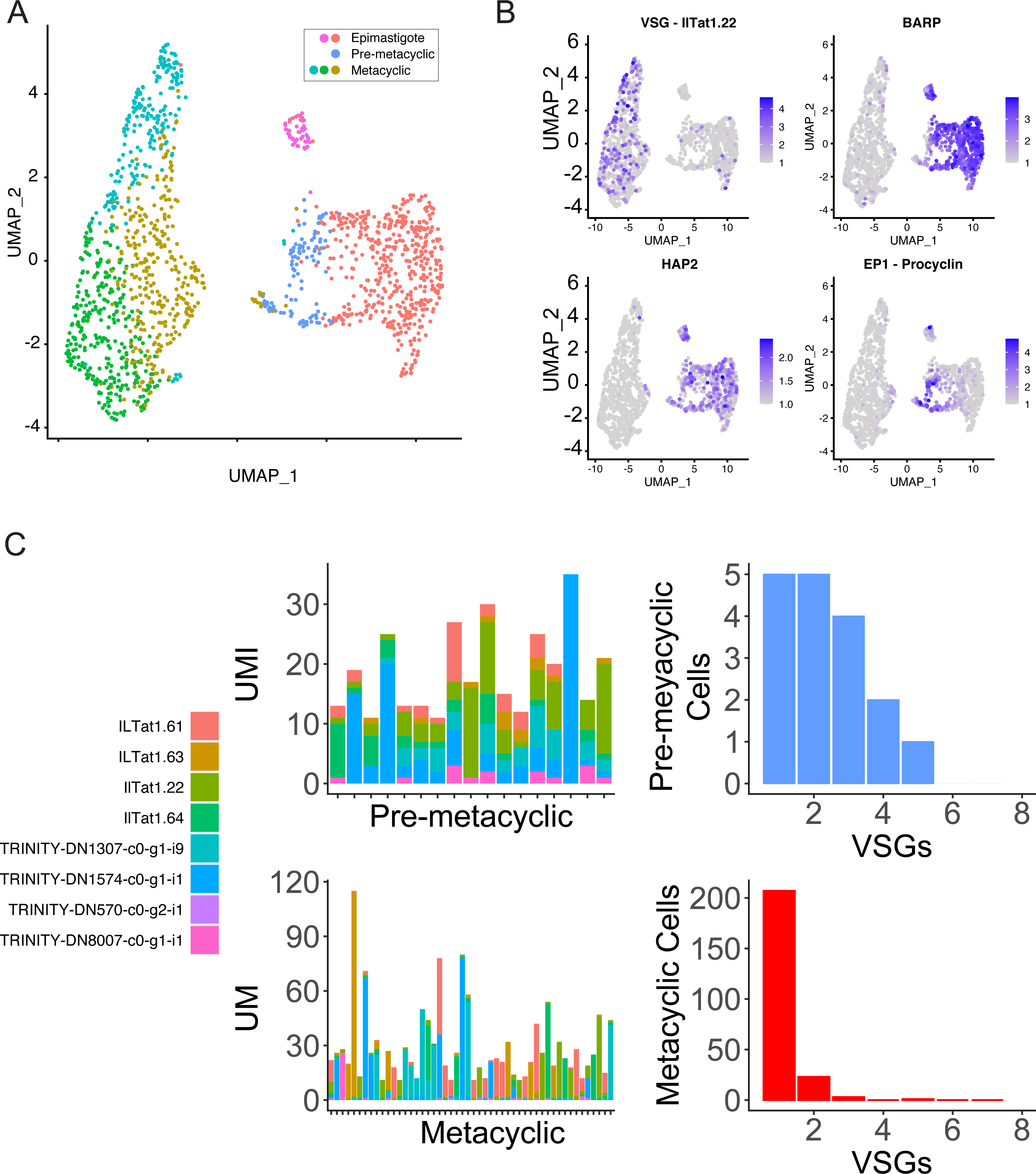
*mVSG* expression analysis of data from Vigneron, O’Neill et al. (2020). UMAP projection of scRNA-seq data indicating the classifications of cells. B. Normalised expression values overlaid on UMAP projections for *VSG* (IlTat1.22 AJ012198.3), *BARP* (Tb927.9.15640), *HAP2* (Tb927.10.10770) and *EP1-Procyclin* (Tb927.10.10260). C. VSG expression profiles for pre-metacyclic (above) and metacyclic cells (below). Left, *VSG* expression data for all pre-metacyclic cells with a total *VSG* UMI count > 10 and a random subset of 50 metacyclic cells with a total *VSG* UMI count > 10. Data are raw (unscaled) UMI counts. Each column represents a cell and each colour a different *VSG*. Right, histogram shows the number of *VSG* expressed for all pre-metacyclic or metacyclic cells (per *VSG* UMI count > 2, all cells in cluster).

## References

Abate, A. R., C. H. Chen, J. J. Agresti and D. A. Weitz (2009). “Beating Poisson encapsulation statistics using close-packed ordering.” Lab Chip 9(18): 2628–2631.

Alibu, V. P., L. Storm, S. Haile, C. Clayton and D. Horn (2005). “A doubly inducible system for RNA interference and rapid RNAi plasmid construction in Trypanosoma brucei.” Mol Biochem Parasitol 139(1): 75–82.

Alsford, S. and D. Horn (2012). “Cell-cycle-regulated control of VSG expression site silencing by histones and histone chaperones ASF1A and CAF-1b in Trypanosoma brucei.” Nucleic Acids Res 40(20): 10150–10160.

Alsford, S., T. Kawahara, L. Glover and D. Horn (2005). “Tagging a T. brucei RRNA locus improves stable transfection efficiency and circumvents inducible expression position effects.” Mol Biochem Parasitol 144(2): 142–148.

Alsford, S., D. J. Turner, S. O. Obado, A. Sanchez-Flores, L. Glover, M. Berriman, C. Hertz-Fowler and D. Horn (2011). “High-throughput phenotyping using parallel sequencing of RNA interference targets in the African trypanosome.” Genome Res 21(6): 915–924.

Aresta-Branco, F., S. Pimenta and L. M. Figueiredo (2016). “A transcription-independent epigenetic mechanism is associated with antigenic switching in Trypanosoma brucei.” Nucleic Acids Res 44(7): 3131–3146.

Batram, C., N. G. Jones, C. J. Janzen, S. M. Markert and M. Engstler (2014). “Expression site attenuation mechanistically links antigenic variation and development in Trypanosoma brucei.” Elife 3: e02324.

Berriman, M., E. Ghedin, C. Hertz-Fowler, G. Blandin, H. Renauld, D. C. Bartholomeu, N. J. Lennard, E. Caler, N. E. Hamlin, B. Haas, U. Bohme, L. Hannick, M. A. Aslett, J. Shallom, L. Marcello, L. Hou, B. Wickstead, U. C. Alsmark, C. Arrowsmith, R. J. Atkin, A. J. Barron, F. Bringaud, K. Brooks, M. Carrington, I. Cherevach, T. J. Chillingworth, C. Churcher, L. N. Clark, C. H. Corton, A. Cronin, R. M. Davies, J. Doggett, A. Djikeng, T. Feldblyum, M. C. Field, A. Fraser, I. Goodhead, Z. Hance, D. Harper, B. R. Harris, H. Hauser, J. Hostetler, A. Ivens, K. Jagels, D. Johnson, J. Johnson, K. Jones, A. X. Kerhornou, H. Koo, N. Larke, S. Landfear, C. Larkin, V. Leech, A. Line, A. Lord, A. Macleod, P. J. Mooney, S. Moule, D. M. Martin, G. W. Morgan, K. Mungall, H. Norbertczak, D. Ormond, G. Pai, C. S. Peacock, J. Peterson, M. A. Quail, E. Rabbinowitsch, M. A. Rajandream, C. Reitter, S. L. Salzberg, M. Sanders, S. Schobel, S. Sharp, M. Simmonds, A. J. Simpson, L. Tallon, C. M. Turner, A. Tait, A. R. Tivey, S. Van Aken, D. Walker, D. Wanless, S. Wang, B. White, O. White, S. Whitehead, J. Woodward, J. Wortman, M. D. Adams, T. M. Embley, K. Gull, E. Ullu, J. D. Barry, A. H. Fairlamb, F. Opperdoes, B. G. Barrell, J. E. Donelson, N. Hall, C. M. Fraser, S. E. Melville and N. M. El-Sayed (2005). “The genome of the African trypanosome Trypanosoma brucei.” Science 309(5733): 416–422.

Brun, R. and Schonenberger (1979). “Cultivation and in vitro cloning or procyclic culture forms of Trypanosoma brucei in a semi-defined medium. Short communication.” Acta Trop 36(3): 289–292.

Butler, A., P. Hoffman, P. Smibert, E. Papalexi and R. Satija (2018). “Integrating single-cell transcriptomic data across different conditions, technologies, and species.” Nat Biotechnol 36(5): 411–420.

Cross, G. A. (1975). “Identification, purification and properties of clone-specific glycoprotein antigens constituting the surface coat of Trypanosoma brucei.” Parasitology 71(3): 393–417.

Cross, G. A. (2017). “http://129.85.245.250/index.html.”

Dalton, R. P., D. B. Lyons and S. Lomvardas (2013). “Co-opting the unfolded protein response to elicit olfactory receptor feedback.” Cell 155(2): 321–332.

Dolezelova, E., M. Kunzova, M. Dejung, M. Levin, B. Panicucci, C. Regnault, C. J. Janzen, M. P. Barrett, F. Butter and A. Zikova (2020). “Cell-based and multi-omics profiling reveals dynamic metabolic repurposing of mitochondria to drive developmental progression of Trypanosoma brucei.” PLoS Biol 18(6): e3000741.

Duraisingh, M. T. and D. Horn (2016). “Epigenetic Regulation of Virulence Gene Expression in Parasitic Protozoa.” Cell Host Microbe 19(5): 629–640.

Edelstein, A. D., M. A. Tsuchida, N. Amodaj, H. Pinkard, R. D. Vale and N. Stuurman (2014). “Advanced methods of microscope control using muManager software.” J Biol Methods 1(2).

Ersfeld, K., R. Docherty, S. Alsford and K. Gull (1996). “A fluorescence in situ hybridisation study of the regulation of histone mRNA levels during the cell cycle of Trypanosoma brucei.” Mol Biochem Parasitol 81(2): 201–209.

Faria, J., L. Glover, S. Hutchinson, C. Boehm, M. C. Field and D. Horn (2019). “Monoallelic expression and epigenetic inheritance sustained by a Trypanosoma brucei variant surface glycoprotein exclusion complex.” Nat Commun 10(1): 3023.

Faria, J., V. Luzak, L. S. M. Muller, B. G. Brink, S. Hutchinson, L. Glover, D. Horn and T. N. Siegel (2021). “Spatial integration of transcription and splicing in a dedicated compartment sustains monogenic antigen expression in African trypanosomes.” Nat Microbiol.

Fedry, J., Y. Liu, G. Pehau-Arnaudet, J. Pei, W. Li, M. A. Tortorici, F. Traincard, A. Meola, G. Bricogne, N. V. Grishin, W. J. Snell, F. A. Rey and T. Krey (2017). “The Ancient Gamete Fusogen HAP2 Is a Eukaryotic Class II Fusion Protein.” Cell 168(5): 904–915 e910.

Figueiredo, L. M., C. J. Janzen and G. A. Cross (2008). “A histone methyltransferase modulates antigenic variation in African trypanosomes.” PLoS Biol 6(7): e161.

Glover, L., S. Hutchinson, S. Alsford and D. Horn (2016). “VEX1 controls the allelic exclusion required for antigenic variation in trypanosomes.” Proc Natl Acad Sci U S A 113(26): 7225–7230.

Grabherr, M. G., B. J. Haas, M. Yassour, J. Z. Levin, D. A. Thompson, I. Amit, X. Adiconis, L. Fan, R. Raychowdhury, Q. Zeng, Z. Chen, E. Mauceli, N. Hacohen, A. Gnirke, N. Rhind, F. di Palma, B. W. Birren, C. Nusbaum, K. Lindblad-Toh, N. Friedman and A. Regev (2011). “Full-length transcriptome assembly from RNA-Seq data without a reference genome.” Nat Biotechnol 29(7): 644–652.

Gunzl, A., T. Bruderer, G. Laufer, B. Schimanski, L. C. Tu, H. M. Chung, P. T. Lee and M. G. Lee (2003). “RNA polymerase I transcribes procyclin genes and variant surface glycoprotein gene expression sites in Trypanosoma brucei.” Eukaryot Cell 2(3): 542–551.

Haanstra, J. R., M. Stewart, V. D. Luu, A. van Tuijl, H. V. Westerhoff, C. Clayton and B. M. Bakker (2008). “Control and regulation of gene expression: quantitative analysis of the expression of phosphoglycerate kinase in bloodstream form Trypanosoma brucei.” J Biol Chem 283(5): 2495–2507.

Hafemeister, C. and R. Satija (2019). “Normalization and variance stabilization of single-cell RNA-seq data using regularized negative binomial regression.” Genome Biol 20(1): 296.

Hanchate, N. K., K. Kondoh, Z. Lu, D. Kuang, X. Ye, X. Qiu, L. Pachter, C. Trapnell and L. B. Buck (2015). “Single-cell transcriptomics reveals receptor transformations during olfactory neurogenesis.” Science 350(6265): 1251–1255.

Hertz-Fowler, C., L. M. Figueiredo, M. A. Quail, M. Becker, A. Jackson, N. Bason, K. Brooks, C. Churcher, S. Fahkro, I. Goodhead, P. Heath, M. Kartvelishvili, K. Mungall, D. Harris, H. Hauser, M. Sanders, D. Saunders, K. Seeger, S. Sharp, J. E. Taylor, D. Walker, B. White, R. Young, G. A. Cross, G. Rudenko, J. D. Barry, E. J. Louis and M. Berriman (2008). “Telomeric expression sites are highly conserved in Trypanosoma brucei.” PLoS One 3(10): e3527.

Hoare, C. A. and F. G. Wallace (1966). “Developmental Stages of Trypanosomatid Flagellates: a New Terminology.” Nature 212(5068): 1385–1386.

Horn, D. (2014). “Antigenic variation in African trypanosomes.” Mol Biochem Parasitol 195(2): 123–129.

International Glossina Genome, I. (2014). “Genome sequence of the tsetse fly (Glossina morsitans): vector of African trypanosomiasis.” Science 344(6182): 380–386.

Islam, S., A. Zeisel, S. Joost, G. La Manno, P. Zajac, M. Kasper, P. Lonnerberg and S. Linnarsson (2014). “Quantitative single-cell RNA-seq with unique molecular identifiers.” Nat Methods 11(2): 163–166.

Janzen, C. J., S. B. Hake, J. E. Lowell and G. A. Cross (2006). “Selective di- or trimethylation of histone H3 lysine 76 by two DOT1 homologs is important for cell cycle regulation in Trypanosoma brucei.” Mol Cell 23(4): 497–507.

Kassem, A., E. Pays and L. Vanhamme (2014). “Transcription is initiated on silent variant surface glycoprotein expression sites despite monoallelic expression in Trypanosoma brucei.” Proc Natl Acad Sci U S A 111(24): 8943–8948.

Kerry, L. E., E. E. Pegg, D. P. Cameron, J. Budzak, G. Poortinga, K. M. Hannan, R. D. Hannan and G. Rudenko (2017). “Selective inhibition of RNA polymerase I transcription as a potential approach to treat African trypanosomiasis.” PLoS Negl Trop Dis 11(3): e0005432.

Klein, A. M., L. Mazutis, I. Akartuna, N. Tallapragada, A. Veres, V. Li, L. Peshkin, D. A. Weitz and M. W. Kirschner (2015). “Droplet barcoding for single-cell transcriptomics applied to embryonic stem cells.” Cell 161(5): 1187–1201.

Klein, G., B. Giovanella, A. Westman, J. S. Stehlin and D. Mumford (1975). “An EBV-genome-negative cell line established from an American Burkitt lymphoma; receptor characteristics. EBV infectibility and permanent conversion into EBV-positive sublines by in vitro infection.” Intervirology 5(6): 319–334.

Kolev, N. G., K. Ramey-Butler, G. A. Cross, E. Ullu and C. Tschudi (2012). “Developmental progression to infectivity in Trypanosoma brucei triggered by an RNA-binding protein.” Science 338(6112): 1352–1353.

Kolev, N. G., E. Ullu and C. Tschudi (2014). “The emerging role of RNA-binding proteins in the life cycle of Trypanosoma brucei.” Cell Microbiol 16(4): 482–489.

L McInnes, J Healy and J. Melville (2018). “Umap: Uniform manifold approximation and projection for dimension reduction.” arxiv arXiv:1802.03426.

Langmead, B. and S. L. Salzberg (2012). “Fast gapped-read alignment with Bowtie 2.” Nature Methods 9(4): 357–359.

Li, H., B. Handsaker, A. Wysoker, T. Fennell, J. Ruan, N. Homer, G. Marth, G. Abecasis, R. Durbin and G. P. D. P. Subgroup (2009). “The Sequence Alignment/Map format and SAMtools.” Bioinformatics 25(16): 2078–2079.

Lyons, D. B., W. E. Allen, T. Goh, L. Tsai, G. Barnea and S. Lomvardas (2013). “An epigenetic trap stabilizes singular olfactory receptor expression.” Cell 154(2): 325–336.

Magklara, A., A. Yen, B. M. Colquitt, E. J. Clowney, W. Allen, E. Markenscoff-Papadimitriou, Z. A. Evans, P. Kheradpour, G. Mountoufaris, C. Carey, G. Barnea, M. Kellis and S. Lomvardas (2011). “An epigenetic signature for monoallelic olfactory receptor expression.” Cell 145(4): 555–570.

Mazutis, L., J. Gilbert, W. L. Ung, D. A. Weitz, A. D. Griffiths and J. A. Heyman (2013). “Single-cell analysis and sorting using droplet-based microfluidics.” Nat Protoc 8(5): 870–891.

Mosser, D. M. and J. F. Roberts (1982). “Trypanosoma brucei: recognition in vitro of two developmental forms by murine macrophages.” Exp Parasitol 54(3): 310–316.

Muller, L. S. M., R. O. Cosentino, K. U. Forstner, J. Guizetti, C. Wedel, N. Kaplan, C. J. Janzen, P. Arampatzi, J. Vogel, S. Steinbiss, T. D. Otto, A. E. Saliba, R. P. Sebra and T. N. Siegel (2018). “Genome organization and DNA accessibility control antigenic variation in trypanosomes.” Nature 563(7729): 121–125.

Navarro, M. and K. Gull (2001). “A pol I transcriptional body associated with VSG mono-allelic expression in Trypanosoma brucei.” Nature 414(6865): 759–763.

Peacock, L., M. Bailey, M. Carrington and W. Gibson (2014). “Meiosis and haploid gametes in the pathogen Trypanosoma brucei.” Curr Biol 24(2): 181–186.

Peacock, L., V. Ferris, R. Sharma, J. Sunter, M. Bailey, M. Carrington and W. Gibson (2011). “Identification of the meiotic life cycle stage of Trypanosoma brucei in the tsetse fly.” Proc Natl Acad Sci U S A 108(9): 3671–3676.

Pinger, J., S. Chowdhury and F. N. Papavasiliou (2017). “Variant surface glycoprotein density defines an immune evasion threshold for African trypanosomes undergoing antigenic variation.” Nature Communications 8(1): 828.

Ramey-Butler, K., E. Ullu, N. G. Kolev and C. Tschudi (2015). “Synchronous expression of individual metacyclic variant surface glycoprotein genes in Trypanosoma brucei.” Mol Biochem Parasitol 200(1-2): 1–4.

Ramirez, F., D. P. Ryan, B. Gruning, V. Bhardwaj, F. Kilpert, A. S. Richter, S. Heyne, F. Dundar and T. Manke (2016). “deepTools2: a next generation web server for deep-sequencing data analysis.” Nucleic Acids Res 44(W1): W160–165.

Roditi, I., M. Carrington and M. Turner (1987). “Expression of a polypeptide containing a dipeptide repeat is confined to the insect stage of Trypanosoma brucei.” Nature 325(6101): 272–274.

Roditi, I., H. Schwarz, T. W. Pearson, R. P. Beecroft, M. K. Liu, J. P. Richardson, H. J. Buhring, J. Pleiss, R. Bulow, R. O. Williams and et al. (1989). “Procyclin gene expression and loss of the variant surface glycoprotein during differentiation of Trypanosoma brucei.” J Cell Biol 108(2): 737–746.

Rotureau, B., I. Subota, J. Buisson and P. Bastin (2012). “A new asymmetric division contributes to the continuous production of infective trypanosomes in the tsetse fly.” Development 139(10): 1842–1850.

Rotureau, B. and J. Van Den Abbeele (2013). “Through the dark continent: African trypanosome development in the tsetse fly.” Front Cell Infect Microbiol 3: 53.

Rueden, C. T., J. Schindelin, M. C. Hiner, B. E. DeZonia, A. E. Walter, E. T. Arena and K. W. Eliceiri (2017). “ImageJ2: ImageJ for the next generation of scientific image data.” BMC Bioinformatics 18(1): 529.

Savage, A. F., N. G. Kolev, J. B. Franklin, A. Vigneron, S. Aksoy and C. Tschudi (2016). “Transcriptome Profiling of Trypanosoma brucei Development in the Tsetse Fly Vector Glossina morsitans.” PLoS One 11(12): e0168877.

Schindelin, J., I. Arganda-Carreras, E. Frise, V. Kaynig, M. Longair, T. Pietzsch, S. Preibisch, C. Rueden, S. Saalfeld, B. Schmid, J. Y. Tinevez, D. J. White, V. Hartenstein, K. Eliceiri, P. Tomancak and A. Cardona (2012). “Fiji: an open-source platform for biological-image analysis.” Nat Methods 9(7): 676–682.

Senabouth, A., S. Andersen, Q. Shi, L. Shi, F. Jiang, W. Zhang, K. Wing, M. Daniszewski, S. W. Lukowski, S. S. C. Hung, Q. Nguyen, L. Fink, A. Beckhouse, A. Pébay, A. W. Hewitt and J. E. Powell (2020). “Comparative performance of the BGI and Illumina sequencing technology for single-cell RNA-sequencing.” NAR Genomics and Bioinformatics 2(2).

Shameer, S., F. J. Logan-Klumpler, F. Vinson, L. Cottret, B. Merlet, F. Achcar, M. Boshart, M. Berriman, R. Breitling, F. Bringaud, P. Bütikofer, A. M. Cattanach, B. Bannerman-Chukualim, D. J. Creek, K. Crouch, H. P. de Koning, H. Denise, C. Ebikeme, A. H. Fairlamb, M. A. Ferguson, M. L. Ginger, C. Hertz-Fowler, E. J. Kerkhoven, P. Mäser, P. A. Michels, A. Nayak, D. W. Nes, D. P. Nolan, C. Olsen, F. Silva-Franco, T. K. Smith, M. C. Taylor, A. G. Tielens, M. D. Urbaniak, J. J. van Hellemond, I. M. Vincent, S. R. Wilkinson, S. Wyllie, F. R. Opperdoes, M. P. Barrett and F. Jourdan (2015). “TrypanoCyc: a community-led biochemical pathways database for Trypanosoma brucei.” Nucleic Acids Res 43(Database issue): D637-644.

Sharma, R., L. Peacock, E. Gluenz, K. Gull, W. Gibson and M. Carrington (2008). “Asymmetric cell division as a route to reduction in cell length and change in cell morphology in trypanosomes.” Protist 159(1): 137–151.

Siegel, T. N., D. R. Hekstra, X. Wang, S. Dewell and G. A. Cross (2010). “Genome-wide analysis of mRNA abundance in two life-cycle stages of Trypanosoma brucei and identification of splicing and polyadenylation sites.” Nucleic Acids Res 38(15): 4946–4957.

Smith, T., A. Heger and I. Sudbery (2017). “UMI-tools: modeling sequencing errors in Unique Molecular Identifiers to improve quantification accuracy.” Genome Res 27(3): 491–499.

Smith, T. K., F. Bringaud, D. P. Nolan and L. M. Figueiredo (2017). “Metabolic reprogramming during the Trypanosoma brucei life cycle.” F1000Res 6.

Stuart, T., A. Butler, P. Hoffman, C. Hafemeister, E. Papalexi, W. M. Mauck, 3rd, Y. Hao, M. Stoeckius, P. Smibert and R. Satija (2019). “Comprehensive Integration of Single-Cell Data.” Cell 177(7): 1888-1902 e1821.

Subota, I., B. Rotureau, T. Blisnick, S. Ngwabyt, M. Durand-Dubief, M. Engstler and P. Bastin (2011). “ALBA proteins are stage regulated during trypanosome development in the tsetse fly and participate in differentiation.” Mol Biol Cell 22(22): 4205–4219.

Tetley, L., C. M. Turner, J. D. Barry, J. S. Crowe and K. Vickerman (1987). “Onset of expression of the variant surface glycoproteins of Trypanosoma brucei in the tsetse fly studied using immunoelectron microscopy.” Journal of Cell Science 87(2): 363–372.

Tetley, L. and K. Vickerman (1985). “Differentiation in Trypanosoma brucei: host-parasite cell junctions and their persistence during acquisition of the variable antigen coat.” J Cell Sci 74: 1–19.

Trenaman, A., L. Glover, S. Hutchinson and D. Horn (2019). “A post-transcriptional respiratome regulon in trypanosomes.” Nucleic Acids Res 47(13): 7063–7077.

Urwyler, S., E. Studer, C. K. Renggli and I. Roditi (2007). “A family of stage-specific alanine-rich proteins on the surface of epimastigote forms of Trypanosoma brucei.” Mol Microbiol 63(1): 218–228.

Van Den Abbeele, J., Y. Claes, D. van Bockstaele, D. Le Ray and M. Coosemans (1999). “Trypanosoma brucei spp. development in the tsetse fly: characterization of the post-mesocyclic stages in the foregut and proboscis.” Parasitology 118 ( **Pt 5**): 469–478.

Vigneron, A., M. B. O’Neill, B. L. Weiss, A. F. Savage, O. C. Campbell, S. Kamhawi, J. G. Valenzuela and S. Aksoy (2020). “Single-cell RNA sequencing of Trypanosoma brucei from tsetse salivary glands unveils metacyclogenesis and identifies potential transmission blocking antigens.” Proc Natl Acad Sci U S A 117(5): 2613–2621.

Yang, X., L. M. Figueiredo, A. Espinal, E. Okubo and B. Li (2009). “RAP1 is essential for silencing telomeric variant surface glycoprotein genes in Trypanosoma brucei.” Cell 137(1): 99–109.

Yates, A. D., P. Achuthan, W. Akanni, J. Allen, J. Allen, J. Alvarez-Jarreta, M. R. Amode, I. M. Armean, A. G. Azov, R. Bennett, J. Bhai, K. Billis, S. Boddu, J. C. Marugan, C. Cummins, C. Davidson, K. Dodiya, R. Fatima, A. Gall, C. G. Giron, L. Gil, T. Grego, L. Haggerty, E. Haskell, T. Hourlier, O. G. Izuogu, S. H. Janacek, T. Juettemann, M. Kay, I. Lavidas, T. Le, D. Lemos, J. G. Martinez, T. Maurel, M. McDowall, A. McMahon, S. Mohanan, B. Moore, M. Nuhn, D. N. Oheh, A. Parker, A. Parton, M. Patricio, M. P. Sakthivel, A. I. Abdul Salam, B. M. Schmitt, H. Schuilenburg, D. Sheppard, M. Sycheva, M. Szuba, K. Taylor, A. Thormann, G. Threadgold, A. Vullo, B. Walts, A. Winterbottom, A. Zadissa, M. Chakiachvili, B. Flint, A. Frankish, S. E. Hunt, I. I. G M. Kostadima, N. Langridge, J. E. Loveland, F. J. Martin, J. Morales, J. M. Mudge, M. Muffato, E. Perry, M. Ruffier, S. J. Trevanion, F. Cunningham, K. L. Howe, D. R. Zerbino and P. Flicek (2020). “Ensembl 2020.” Nucleic Acids Res 48(D1): D682–D688.

Zhang, X., T. Li, F. Liu, Y. Chen, J. Yao, Z. Li, Y. Huang and J. Wang (2019). “Comparative Analysis of Droplet-Based Ultra-High-Throughput Single-Cell RNA-Seq Systems.” Mol Cell 73(1): 130–142 e135.

Zilionis, R., J. Nainys, A. Veres, V. Savova, D. Zemmour, A. M. Klein and L. Mazutis (2017). “Single-cell barcoding and sequencing using droplet microfluidics.” Nat Protoc 12(1): 44–73.

Zomerdijk, J. C., R. Kieft and P. Borst (1991). “Efficient production of functional mRNA mediated by RNA polymerase I in Trypanosoma brucei.” Nature 353(6346): 772–775.

